# TRAIL signalling promotes entosis in colorectal cancer

**DOI:** 10.1101/2020.09.29.315317

**Authors:** Emir Bozkurt, Heiko Düssmann, Manuela Salvucci, Brenton L. Cavanagh, Sandra Van Schaeybroeck, Daniel B. Longley, Seamus J. Martin, Jochen H. M. Prehn

**Affiliations:** Department of Physiology and Medical Physics, Centre for Systems Medicine, Royal College of Surgeons in Ireland, 123 St. Stephen’s Green, Dublin 2, Ireland; Department of Genetics and Bioengineering, Faculty of Engineering, Izmir University of Economics, Balcova, Izmir, Turkey; Cellular and Molecular Imaging Core, Royal College of Surgeons in Ireland, D02 YN77 Dublin 2, Ireland; Centre for Cancer Research and Cell Biology, School of Medicine, Dentistry and Biomedical Sciences, Queen’s University Belfast, 97 Lisburn Road, Belfast, BT9 7BL, UK; Molecular Cell Biology Laboratory, Department of Genetics, The Smurfit Institute, Trinity College, Dublin, Ireland

**Keywords:** TRAIL, entosis, cell-in-cell structures, apoptosis, caspase-8, colorectal cancer

## Abstract

Entosis is a form of non-phagocytic cell-in-cell (CIC) interaction where a living cell enters into another. Tumours show evidence of entosis, however factors controlling entosis remain to be elucidated. Here we find that the death receptor ligand TRAIL is a potent activator of entosis in colon cancer cells. CLEM/3D confocal microscopy analysis revealed ultrastructural features of entosis and subsequent entotic cell death of inner cells upon TRAIL treatment. Induction of entosis and apoptosis by TRAIL were mutually exclusive events but both required the presence of caspase-8. *Bax/Bak* double knock-out or caspase inhibition altered the fate of inner cells from entotic cell death to survival and escape. Analysis of colorectal cancer tumours showed a significant association between expression levels of TRAIL and CICs. Notably, the presence of CICs in the invasive front regions of colorectal tumours was significantly correlated with adverse patient prognosis.

## Introduction

Cell death is essential for maintenance of homeostasis in multicellular organisms. Studies over the last two decades have revealed that cell death can occur via multiple pathways, many of which are highly interconnected and may occur simultaneously. Apoptosis is a form of controlled cell death associated with distinct morphological changes such as membrane blebbing and nuclear condensation and fragmentation^1^. Apoptosis is triggered through a family of cysteine proteases known as caspases. Depending on how caspases are activated, apoptosis can be induced by the intrinsic and extrinsic pathways. The intrinsic pathway involves permeabilisation of the mitochondrial outer membrane (MOMP) by Bax and Bak, resulting in the release of mitochondrial intermembrane proteins facilitating activation of caspase-9 and executioner caspases −3, −6, −7. Initiation of the extrinsic pathway requires binding of specific death ligands to transmembrane receptors to initiate the formation of death-inducing signalling complex (DISC) ultimately resulting in caspase-8 activation. Caspase-8 can then trigger a downstream apoptotic cascade either by directly activating the effector caspases or engaging with the mitochondrial pathway by cleaving BID, which leads to MOMP and further activation of executioner caspases^2^

Tumor necrosis factor-related apoptosis-inducing ligand (TRAIL) is a type II transmembrane protein mainly expressed on the surface of a variety of normal cells^3^ as well as most immune cells^4,5^. Both membrane-bound and soluble TRAIL can bind to its death domain (DD) containing receptors (TRAIL-R1/DR4 and TRAIL-R2/DR5) on cancer cells to initiate apoptosis by activating the extrinsic pathway^5^. Because DR4 and DR5 are frequently overexpressed in many cancers compared to normal tissues, targeting these receptors therapeutically has attracted much interest^6^. However, 1^st^ generation agents failed in clinical trials leading to the need to re-evaluate TRAIL signalling from different perspectives, one of which focuses on understanding the nature of cancer cells that survive in response to TRAIL treatment^7^. When cancer cells are exposed to TRAIL, even at high TRAIL levels assumed to saturate all of its receptors, only a fraction of cells undergo complete cell death^8^. Accumulating evidence suggests that this behaviour arises from the parallel activation of cell survival pathways upon TRAIL treatment, as cell death can be augmented by translation inhibitors such as cycloheximide (CHX)^9^. Indeed, although activation of TRAIL signalling recruits proteins such as FADD for apoptosis induction, binding of other regulators such as cellular FLICE-like inhibitory protein (c-FLIP) can switch the pathway towards the activation of survival signalling and generation of cytokines that may promote tumour growth^10,11^.

The phenomenon of cell-in-cell structures (CICs) have been noticed for over a century^12^, however, it was only very recently discovered that not all CICs are formed by phagocytosis-like activity. Entosis is a homotypic cell-in-cell invasion process, by which one living cell (‘inner cell’) actively enters into another (non-phagocytic) living cell (‘outer cell’) of the same type in a Rho/ROCK signalling-dependent manner^13^.

Inner cells are initially alive after invaded into another cell and can even undergo cell division inside the outer cell; moreover, they can also be released. However, the vast majority of inner cells undergo degradation inside outer cells by an autophagic lipidation-dependent lysosomal cell death mechanism known as entotic cell death^14^. Remarkably, even though entotic cell death can result in the elimination of cells at the individual level, entosis can confer a survival advantage to the outer cells under stress conditions^15,16^. The presence of *TP53*^17^ and *KRAS*^16^ mutations increase the likelihood of cancer cells undergoing entosis. Several human malignancies including lung, head and neck, breast, colon, stomach, liver, cervical cancer and melanoma show evidence of CICs^13,17–19^; however, the clinical relevance of CICs remains to be elucidated.

Here, while analysing cell-to-cell variability during TRAIL-mediated caspase activation of colorectal cancer cells at the single-cell level during time-lapse imaging, we discovered that TRAIL activates both apoptotic and entotic cell death in colon cancer cells. We demonstrate that TRAIL-induced entosis requires caspase-8, and that inhibition of apoptosis signalling downstream of caspase-8 significantly altered cell fate during TRAIL-induced entosis. Furthermore, we demonstrate that the presence of CICs in the invading margins of tumours from colorectal cancer patients correlates with key components of the TRAIL signalling pathway and is an independent prognostic marker of disease recurrence for stage 3 colorectal cancer.

## Results

### Single-cell time-lapse microscopy reveals simultaneous induction of entosis and apoptosis in HCT116 colon cancer cells upon TRAIL stimulation

We measured the kinetics of caspase activation by single-cell time-lapse microscopy in HCT116 colon cancer cells stably expressing a Cyan Fluorescent Protein (CFP)-IETD-Venus fluorescent protein (Venus) FRET probe, a fluorescent-based caspase activation reporter that can detect caspase-8-like and caspase-3-like activity during activation of the extrinsic apoptosis pathway^20^ (Figure 1A, 1B). Cells were also incubated with TMRM to indicate the time of mitochondrial outer membrane permeabilization (MOMP) during apoptosis. We quantified the changes in CFP/FRET emission ratio in control cells as well as in response to TRAIL with or without CHX. Typically, cells responding to TRAIL/CHX exhibit a moderate increase in CFP/FRET emission ratio indicating activation of caspase-8, followed by a drop in TMRM intensity as an indicator of MOMP, then a sharp increase in CFP/FRET ratio indicating the activation of caspase-3 and other executioner caspases^8,20^ (Figure 1A). We did not observe any changes in CFP/FRET emission ratio in control cells (not shown). In line with previous reports^8,21^, HCT116 cells exhibited a heterogeneous response in IETD substrate cleavage in response to TRAIL and in the absence of CHX. A proportion of cells showed IETD substrate cleavage followed by loss of TMRM intensity, ultimately leading to membrane blebbing and cell shrinkage again indicating that apoptosis was induced. Other cells exhibited a slow substrate cleavage, retained intact morphology and TMRM staining suggesting that these cells survived TRAIL treatment (Figure 1C-left, 1E). Overall, the omission of the protein synthesis inhibitor CHX yielded a much more heterogeneous IETD substrate cleavage when compared to TRAIL plus CHX treatment (Figure 1C). To quantify the difference in IETD substrate cleavage kinetics, we fitted single-cell traces to a Boltzmann sigmoid equation and calculated the time to reach 50% substrate cleavage. Presence of CHX significantly reduced the time to reach 50% substrate cleavage and showed a lower standard deviation (Figure 1D).

**Figure 1.**
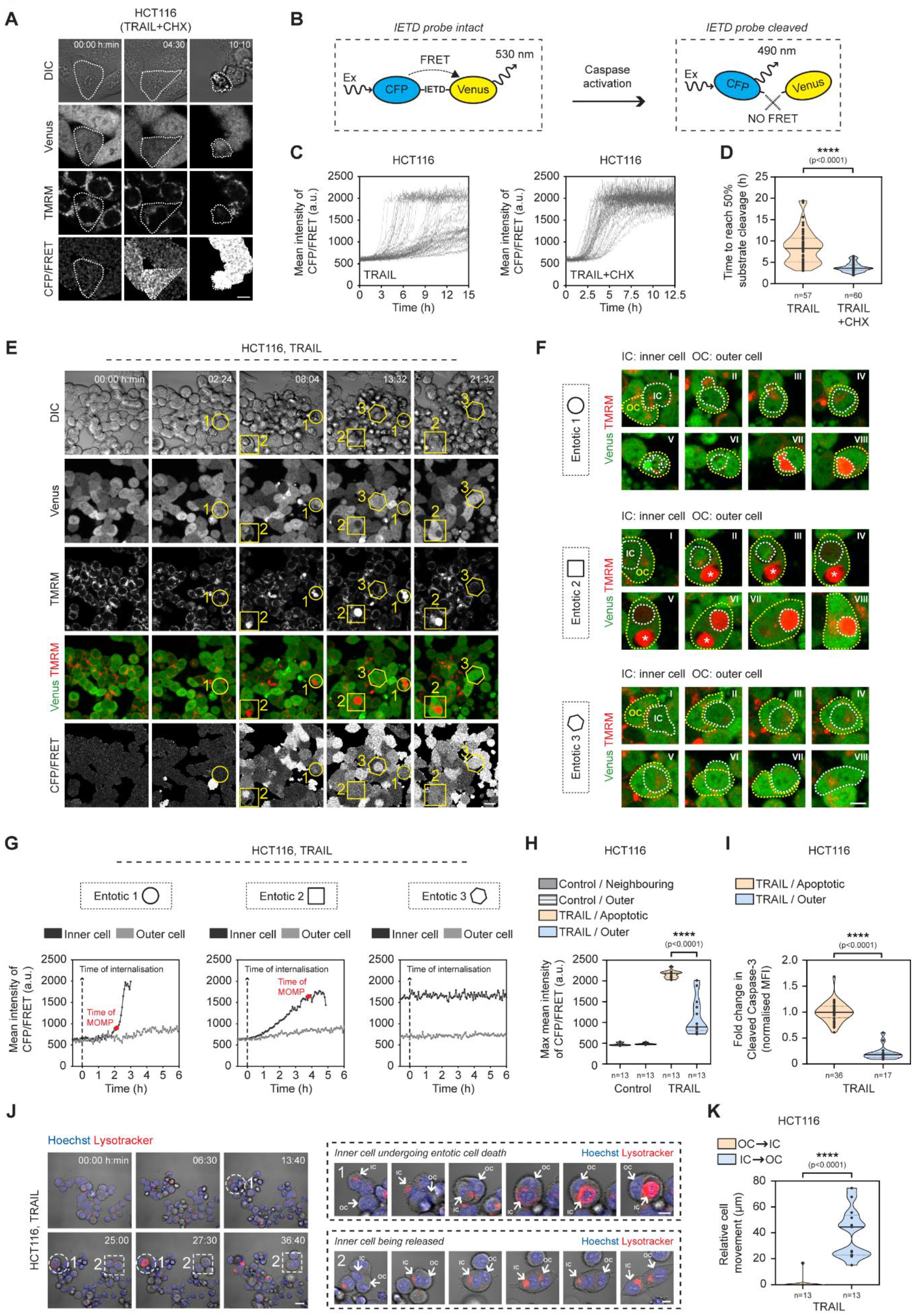
Single-cell time-lapse microscopy reveals simultaneous induction of entosis and apoptosis upon TRAIL stimulation in HCT116 cells. **A-D.** HCT116 cells exhibit heterogeneity in caspase activation kinetics in response to TRAIL but not TRAIL+CHX **A**. Representative time-lapse images of HCT116 cells stably expressing IETD-FRET probe treated with TRAIL+CHX. Venus, TMRM and CFP/FRET emission ratio images are shown. Depicted area highlights an apoptotic cell. Scale bar: 20 μm. Images are representatives from three experiments. **B.** Schematic representation of the intact (left) and the cleaved (right) forms of IETD-FRET probe. **C.** Quantification of single-cell CFP/FRET emission ratio traces in response to TRAIL (left) and TRAIL+CHX (right) (n=58 and n=60 cells were quantified from three experiments). **D.** Single-cell IETD substrate cleavage kinetics (time to reach 50% substrate cleavage) in response to TRAIL and TRAIL+CHX (data obtained by fitting single-cell traces to Boltzmann sigmoid equation). Data are shown as individual values for each cell as well as median and quartiles, n=3, statistical significance was tested using unpaired two tailed t-test, ****p<0.0001. **E-F.** Live-cell TMRM staining accumulates in inner cells during entotic cell death. **E.** Representative time-lapse images of HCT116 IETD cells treated with TRAIL showing formation of apoptotic and entotic structures. DIC, Venus, TMRM and CFP/FRET emission ratio images are shown. Venus and TMRM images are combined and colour-coded as green and red, respectively. Representative entosis events in are highlighted in a circle (entotic 1), a square (entotic 2) and a hexagon (entotic 3). Scale bar: 20 μm. Images are representatives from three experiments. Combined Venus and TMRM images highlighting stages of depicted entosis events. White-dashed and yellow-dashed lines indicate inner cells (IC) and outer cells (OC), respectively. Asterisk indicates another entotic event within in an outer cell of interest. Entotic1: *inner cell shows apoptotic features, then undergoes entotic cell death;* Entotic2: *inner cell shows caspase activation and loss of TMRM signal, then undergoes entotic cell death;* Entotic3: *inner cell with caspase activation invades into another cell, then is released.* Scale bar: 20 μm. **G-I.** Outer cells exhibit slow caspase activation kinetics and show less Cleaved Caspase-3 levels in TRAIL treatment. **G.** Quantification of CFP/FRET emission ratio traces of inner (black) and outer (grey) cells in entotic 1 (circle), entotic 2 (square) and entotic 3 (hexagon). Dashed line with arrow indicates time of internalisation. Red dot on the single cell trace of interest represents time of TMRM intensity loss. Quantification of maximum mean values of CFP/FRET emission ratio traces in outer cells, apoptotic cells or neighbouring cells in HCT116 treated with or without TRAIL. Single-cell CFP/FRET emission ratio traces of n=52 cells (13 cells per group) were generated over 20 hours of time-lapse microscopy and maximum value in each trace was recorded. Data are shown as individual values for each cell as well as median and quartiles from three experiments. Statistical significance was tested using one-way ANOVA followed by Tukey’s multiple comparison test. ****p<0.0001, ns: not significant. **I.** Quantification of immunofluorescence staining of Cleaved Caspase-3 (Asp175) levels in apoptotic cells (n=36) and outer cells (n=30) in HCT116 treated with TRAIL. Entosis events were characterised using 3D confocal images of DIC and Hoechst by detecting a round-shaped Hoechst-stained cell inside a vacuolar structure within another Hoechst-stained cell showing a crescent-shaped nuclear morphology. Cells showing fragmented nuclei and increased Cleaved Caspase-3 intensity were considered as apoptotic. Data are shown as individual values for each cell as well median and quartiles from three experiments, statistical significance was tested using unpaired two tailed t-test, ****p<0.0001. **J-K.** Long-term time-lapse microscopy verifies induction of entotic structures by TRAIL treatment. Representative time-lapse images of DNA (Hoechst) and lysosome (Lysotracker) staining in HCT116 cells showing formation of entotic structures in TRAIL treatment. Representative entosis events of an inner cell undergoing entotic cell death (1) and an inner cell is being released (2) are highlighted in a circle with dashed line and a square with dashed line (left). Formation and stages of depicted entosis events are shown (right). Hoechst and Lysotracker staining are colour-coded as blue and red, respectively. Inner cells (IC) and outer cells (OC) are highlighted with white arrows. **K.** Quantification of relative cell movement of inner and outer cells in TRAIL-treated HCT116 cells. n=13 entotic structures from three experiments were monitored by time-lapse microscopy. Data are shown as individual values for each cell as well as median and quartiles, statistical significance was tested using unpaired two tailed t-test, ****p<0.0001.

While investigating cell-to-cell variability in caspase activation dynamics upon TRAIL stimulation, we noticed that a proportion of cells entered into neighbouring cells and appeared inside a large vacuolar structure (Figure 1E, 1F, Supplementary Video (SV) 1). We further noted that, before cell-in-cell invasion, inner cells showed an active movement towards outer cells. In separate time-lapse experiments, vacuolar structures showed Lysotracker accumulation and inner cells in such structures underwent degradation indicating that inner cells were degraded inside of acidic structures (Figure 1J, 1K, SV2, SV3). Of note, Lysotracker has previously been used to monitor entotic cell death^13^. We also observed occasions of not only single cells but multiple cells invading into their neighbours (Figure 1E, 1F). A period of time after cell-in-cell invasion, the majority of inner cells also showed a reduction in the intensity of Venus, which is quenched at acidic pH^22^. Remarkably, in coincidence with the reduction in Venus intensity, these cells exhibited a significant increase in the TMRM signal (Figure 1E, 1F, Entotic 1 and Entotic 2). To further examine whether this increase was associated with inner cell death, we performed time-lapse microscopy to simultaneously measure fluorescence intensities of Venus, TMRM, and Lysotracker. We found that the reduction in Venus and the increase in TMRM fluorescence coincided with the accumulation of Lysotracker in inner cells that underwent degradation (Figure S1A, SV4). Treatment with FCCP, a protonophore that dissipates the mitochondrial membrane potential showed an immediate drop in TMRM intensity in outer and neighbouring cells, however, inner cells undergoing degradation did not show a drop in TMRM fluorescence intensity, suggesting that the increase in TMRM fluorescence intensity was not related to changes in the mitochondrial membrane potential (Figure S1B). Interestingly, reduction in Venus, accumulation of TMRM and Lysotracker occurred only when inner cells underwent degradation. In rare cases, inner cells were released hours after cell-in-cell invasion (Figure 1F, Entotic 3). There was no change in Venus, TMRM and Lysotracker fluorescence intensities when inner cells were released (Figure S1A).

To examine the associations among entosis, entotic cell death and IETD substrate cleavage upon TRAIL stimulation, we quantified CFP/FRET emission ratios focusing on cell-in-cell structures. As shown in Figure 1F and 1G (Entotic 1), while appearing inside another cell, inner cells exhibited an increase in IETD substrate cleavage and a drop in TMRM intensity similar to what we observed in classical apoptotic cells. Later on, these cells showed a reduction in Venus fluorescence and accumulation of TMRM indicating that they underwent entotic cell death (Figure 1G, Entotic 1, Entotic 2). In some cases, we also observed that cells showing IETD substrate cleavage entered into neighbouring cells but were released hours after internalisation (Figure 1G, Entotic 3). We also monitored spontaneous entotic events in control cells that were not exposed to TRAIL. Inner cells did not display any change in IETD substrate cleavage before entering, or while inside outer cells during spontaneous entosis. However, reduction in Venus fluorescence intensity and TMRM fluorescence accumulation were still evident at later entotic stages. This suggested that IETD substrate cleavage in inner cells during TRAIL-induced entosis was specifically related to this form of entosis.

We next focused on whether IETD substrate cleavage changed in outer cells upon TRAIL stimulation. Remarkably, outer cells showed a very slow IETD substrate cleavage similar to TRAIL treatment surviving cells (Figure 1G, Entotic 1,2,3). To examine whether outer cells showed less substrate cleavage compared to neighbouring apoptotic cells, we compared the maximum value of CFP/FRET emission ratio kinetics in outer cells in controls and 24 hours TRAIL-treated cells. We found that maximum IETD substrate cleavage values were significantly lower in outer cells compared to neighbouring apoptotic cells. We observed no difference in CFP/FRET emission ratio when we compared outer cells with neighbouring cells in untreated cultures (Figure 1H).

We next performed immunofluorescence staining and confocal microscopy analysis for cleaved caspase-3 in conjunction with nuclear staining (Hoechst) in cells treated with or without TRAIL. As expected, TRAIL treatment resulted in a significant increase in cleaved caspase-3 intensity in all treated cells compared to control cells (Figure S2A). In parallel, we observed that most inner cells (83.57 ± 5.78 %) showed an increase in cleaved caspase-3 staining (Figure S2B). Moreover, we could occasionally observe fragmented nuclear morphology as well as the release of cytochrome c in a fraction of inner cells indicating that these cells indeed showed to some degree ‘apoptotic features’ during entotic cell death. However, the majority of inner cells did not show apoptotic features and exhibited foci formation for both cleaved caspase-3 and cytochrome c during entotic cell death (Figure S2B and S2C). As expected, we found that outer cells showed significantly lower levels of cleaved caspase-3 staining than neighbouring apoptotic cells (Figure 1I). To examine the long-term survival of outer cells, we monitored 42 entotic structures by time-lapse microscopy in HCT116 cells treated with TRAIL for 48 hours. Based on cell morphology, TMRM signal and Venus signal in outer cells, we found that 38 out of 42 outer cells survived in TRAIL treatment. These results suggested that outer cells gained a survival advantage due to entosis.

### Large-scale HCS-based analysis validates simultaneous induction of entosis and apoptosis in TRAIL but not TRAIL/CHX treated cells

Although entosis has a unique cytological structure making the events possible to identify, we realised that there was a need to develop an unbiased method of quantifying entosis. To address this, we sought to design an automated, large-scale analysis for simultaneous detection of entotic and cell death events by using a high-content screening platform (HCS) (Figure 2). Briefly, cells that have been seeded into 96 well plates were labelled with Hoechst, Propidium iodide and Lysotracker and exposed to different treatments (Figure 2A). After recording 12 random fields per well for 72 hours, images were segmented using CellProfiler v2.2 and combined to detect entotic events (Figure 2B, C). We were able to successfully detect entotic events based on the accumulation of Lysotracker staining that co-localised with Hoechst but not PI staining in inner cells (Figure 2D, arrows). Moreover, by combining brightfield, Hoechst and Lysotracker images, we observed cells inside a large vacuole within another cell without showing Lysotracker accumulation or PI uptake, suggesting that they were early entotic events (Figure 2D, dashed arrows). Cells that showed PI uptake were considered as dead (PI^+^ cells) (see Materials and Methods for further details). TRAIL significantly increased the number of PI-positive cells as compared to control. Moreover, the presence of CHX further enhanced TRAIL-induced cell death (Figure 2E). Strikingly, TRAIL resulted in a significant increase in entosis events in a time-dependent manner, as compared to control cells. In contrast, we observed a reduction in entosis events in the presence of CHX (Figure 2F). Importantly, different cell line models (colon: LS180; breast: MCF7) (Figure S3A and B), as well as spheroids prepared from HCT116 cells, likewise showed significant increases in entosis events in response to TRAIL treatment (Figure S3C, SV5).

**Figure 2.**
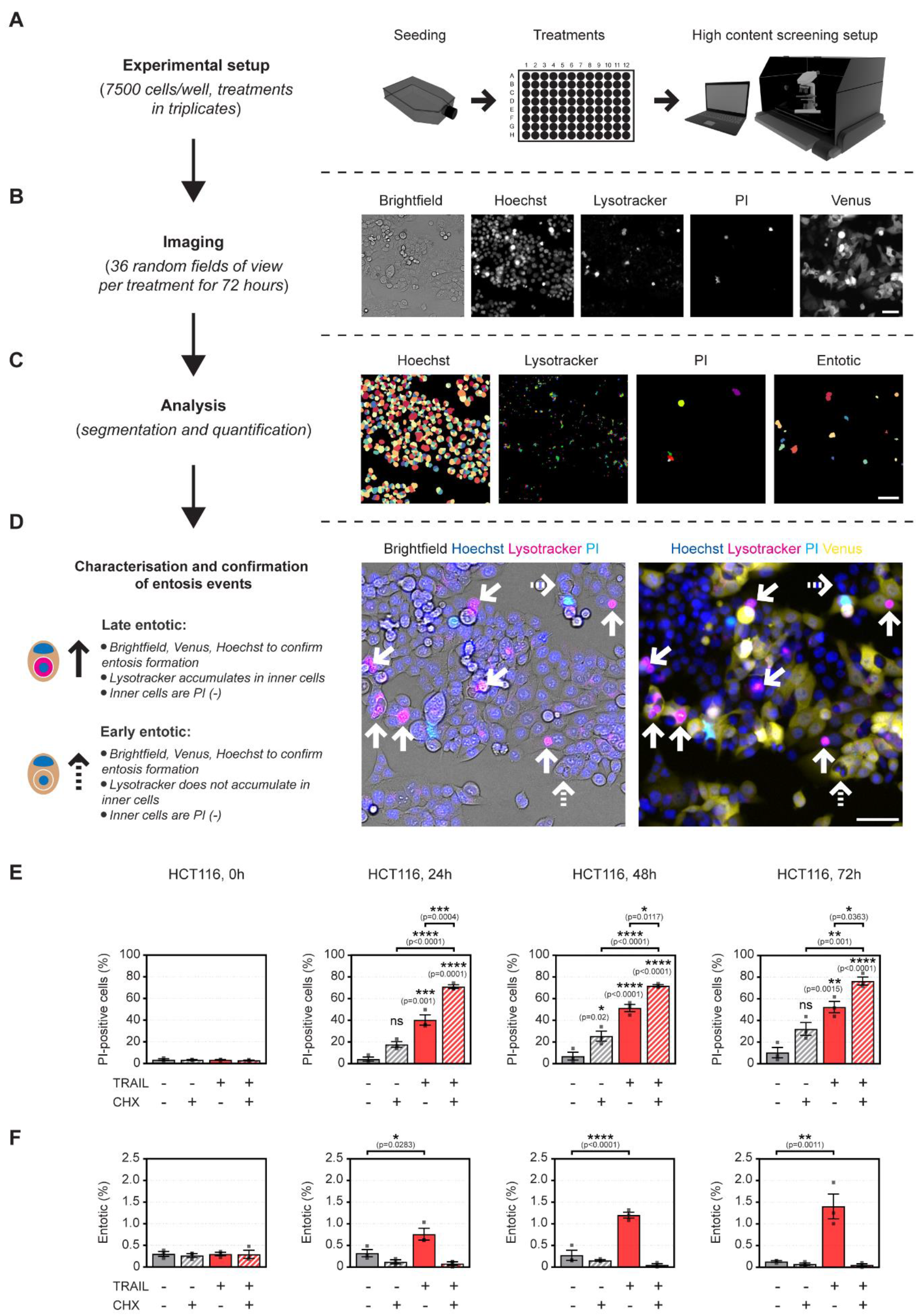
Large-scale HCS-based quantification of simultaneous induction of entosis and apoptosis upon TRAIL and TRAIL+CHX stimulation. **A-D.** Overview of steps involved in analysing entotic structures by HCS. **A.** After seeding (7500 cells/well), staining (Hoechst, Lysotracker, PI) and treatment, (**B**) 12 random fields of view per well were recorded for 72 hours. **C.** CellProfiler defined cell nuclei, lysosomes and PI-positive cells, then overlap of Hoechst and Lysotracker was used to detect late stage entosis events. **D.** Entosis events detected by CellProfiler were manually confirmed using overlaid images of Brightfield, Hoechst, Lysotracker, PI and Venus by analysing inner cells showing accumulation of Lysotracker that co-localises with Hoechst but not PI within another Hoechst-stained cell showing a crescent-shaped nuclear morphology (Late entotic, indicated with arrows). Early stage entosis events were detected manually by analysing round-shaped Hoechst stained cells inside a large vacuole within another Hoechst-stained cell showing a crescent-shaped nuclear morphology (Early entotic, indicated with dashed arrows). Hoechst, Lysotracker, PI and Venus images are colour-coded as blue, magenta, cyan and yellow, respectively. Images are representatives from three experiments. Scale bars: 50 μm. **E-F.** TRAIL but not TRAIL+CHX promotes entosis in HCT116 cells. Quantification of PI positive cells. **(E)**and entotic structures **(F)**in HCT116 cells treated with or without CHX, TRAIL and TRAIL+CHX for 72 hours. Data are shown as individual values for each experiment as well as means ± S.E.M., n=3, statistical significance was tested using one-way ANOVA followed by Tukey’s multiple comparison test. Asterisks on top of individual bars indicate comparisons with control. *p<0.05, **p<0.01, ***p<0.001, ****p<0.0001, ns: not significant.

### Characterisation of entotic ultrastructures and features of entotic cell death induced by TRAIL treatment

Next, we performed a set of experiments for in-depth characterisation of entosis events during TRAIL treatment. First, we analysed the morphological changes in early and late-stage entotic structures in control-treated and TRAIL-treated cells by staining plasma membrane (β-catenin), nuclei (Hoechst) and lysosomal accumulation (Lysotracker). As shown in Figure 3A, 3D confocal imaging of early and late-stage entosis events verified that inner cells were fully internalised. While β-catenin staining was detectable around inner cells at early stages (Figure 3A, top), we did not detect β-catenin staining in inner cells at late stages (Figure 3A, bottom), suggesting that β-catenin underwent degradation in inner cells during entotic cell death. We did not observe morphological difference in plasma membranes, nuclei and lysosomal accumulation when we compared entotic structures in controls with the ones in TRAIL-treated cells (not shown). To confirm whether late-stage inner cells underwent lysosomal cell death in TRAIL treatment, we performed 3D confocal microscopy analysis to measure cleavage of a specific fluorogenic Cathepsin B substrate (Z-Arg-Arg-AMC) in TRAIL-treated cells. As shown in Figure 3B, all inner cells that showed Lysotracker accumulation also showed a significant increase in Cathepsin B activity (Figure 3C) suggesting that late-stage entotic cells underwent Cathepsin-B-mediated lysosomal cell death in TRAIL treatment.

**Figure 3.**
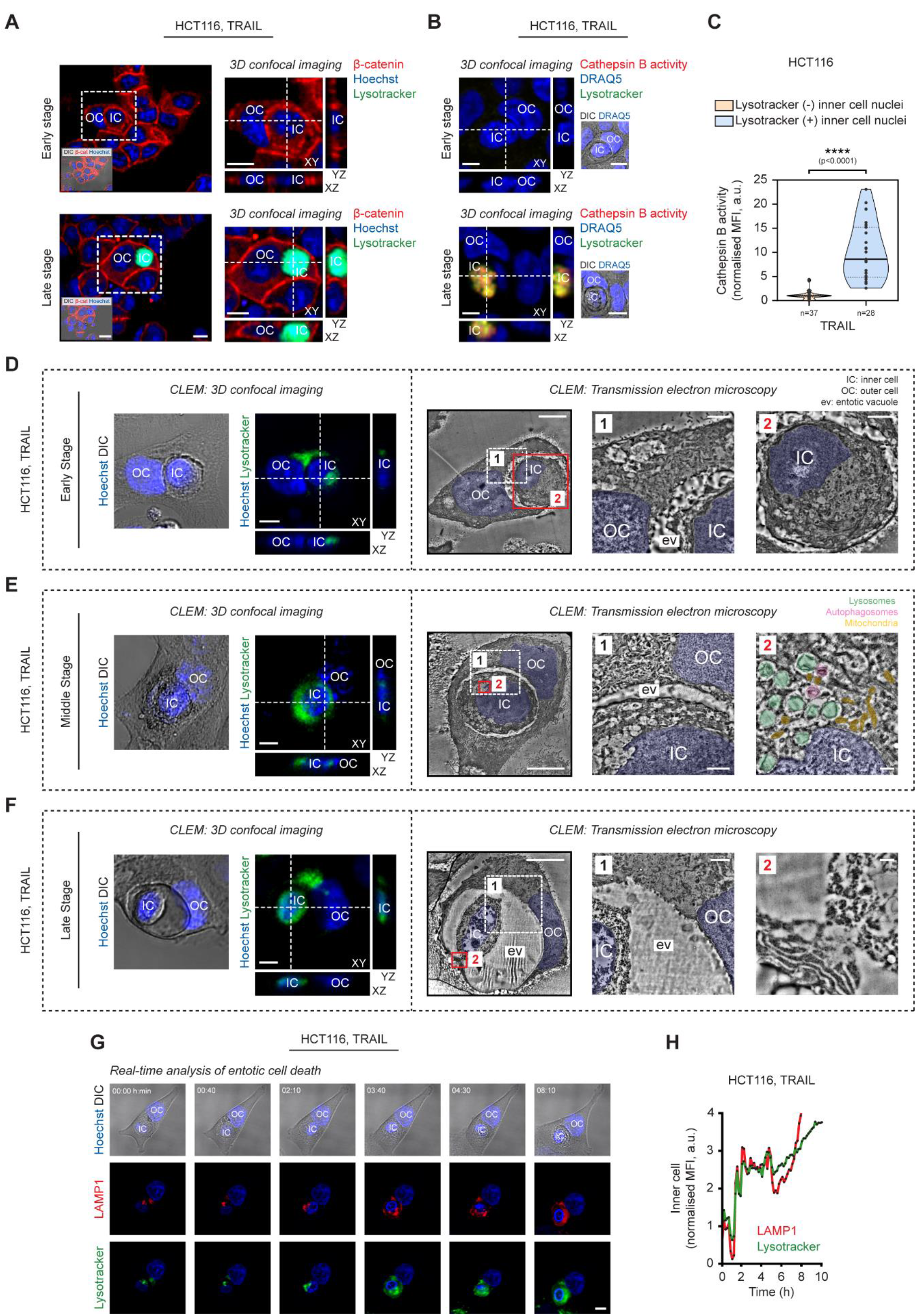
3D Confocal microscopy and CLEM analysis of structural features of entosis in TRAIL-stimulated HCT116 cells. **A-C.** Late stage inner cells undergo Cathepsin B-mediated lysosomal cell death. **A.** 3D confocal microscopy imaging of β-catenin (red), Hoechst (blue) and Lysotracker (green) confirms complete cell internalisation in TRAIL-treated cells. Representative confocal images of early (top) and late (bottom) stage entotic structures are shown. Orthogonal views through z-stacks of depicted areas are shown next to each image. OC: outer cell, IC: inner cell, Scale bars: 10 μm. Images are representatives from three experiments. **B.**Representative 3D confocal microscopy images of fluorogenic Cathepsin B substrate (red), DRAQ5 (Blue) and Lysotracker (green) of early and late stage entotic cells are shown. IC: inner cell, OC: outer cell, Scale bars: 10 μm. Images are representatives from three experiments. Quantification of Cathepsin B activity in early (Lysotracker negative inner cell nuclei) and late (Lysotracker positive inner cell nuclei) stage entotic cells. n=37 (early stage) and n=28 (late stage) inner cells were quantified from three experiments. Data are shown as individual values for each cell as well as median and quartiles, statistical significance was tested using unpaired two tailed t-test, ****p<0.0001. **D-F.** Correlative Light and Electron Microscopy (CLEM) analysis of entotic ultrastructures in TRAIL-treated cells. Representative 3D confocal microscopy images of Hoechst (blue) and Lysotracker (green) staining of early (**D**), middle (**E**) and late (**F**) stages of entosis events are shown (left, Scale bars: 10 μm). TEM images of each corresponding entosis event are presented (right). Two depicted areas (white and red squares) imaged in higher magnifications are shown next to each image. Green, magenta and yellow pseudo-coloured areas indicate lysosomes, autophagosomes and mitochondria, respectively. Blue pseudo-coloured areas indicate nuclei. IC: inner cell, OC: outer cell, ev: entotic vacuole. Scale bars in TEM images, from left to right: 10 μm, 2 μm and 1 μm (**a**) or 500 nm (**b**and **c**). Images are representatives from three experiments. **G-I.** Real-time analysis of lysosomal events during entotic cell death in TRAIL-treated cells. **G.** Representative time-lapse images of Lysotracker staining (green) in HCT116 cells stably expressing Lamp1-Scarlet plasmid (red) showing an inner cell undergoing entotic cell death, scale bar: 10 μm, OC: outer cell, IC: inner cell. Images are representatives from three experiments. **H.** Quantification of LAMP1 (left) and Lysotracker (right) mean fluorescence intensities in depicted inner cell undergoing entotic cell death.

To examine the cellular ultrastructures of TRAIL-induced entosis in more detail, we performed Correlative Light and Electron Microscopy (CLEM) analysis in TRAIL-treated HCT116 cells. First, we used 3D confocal microscopy to image Hoechst and Lysotracker staining in different stages of entosis. After verifying that inner cells were fully internalised, we performed TEM analysis to observe the ultrastructures in the corresponding entosis events (Figure 3D-F). As expected, at all stages, inner cells showed a round-shaped morphology within outer cells, an entotic vacuole between inner and outer cells was visible, and the nucleus of outer cells was pushed towards the periphery forming a crescent-like shape. During the early stages, both inner and outer cells retained cell membranes, preserved cytoplasmic organelles, and exhibited an intact nuclear membrane, nucleoli and dispersed chromatin (Figure 3D). During middle stages, we observed an increase in lysosomal staining as well as accumulation of several lysosomes, autophagosomes and autophagolysosomes in inner cells (Figure 3E). We were able to further confirm autophagosome structures in the corresponding inner cell by immunogold labelling for LC3 (Figure S4). Moreover, some mitochondria juxtaposed to lysosomes, autophagosomes and/or autophagolysosomes, but were not localised inside of them. Interestingly, when we looked at the nucleus of inner cells, we observed that some lysosomes and autophagosomes were localised in the areas where the nuclear membrane was disrupted (Figure 3E). During the late stages, inner cells showed an increase in lysosomal staining co-localising with nuclear staining. Furthermore, TEM analysis showed that both cytoplasmic and nuclear content of inner cells exhibited a complete degradation pattern. We could observe the remnants of inner cells; however, none of the cytoplasmic organelles was detectable (Figure 3F). Although some inner cells displayed apoptotic nuclear morphology, most inner cells did not show an apoptosis-like nuclear pattern.

To fully explore the stages of entotic cell death during TRAIL treatment, we performed time-lapse microscopy in HCT116 cells transfected with Lysosomal-associated membrane protein 1 (LAMP1)-mScarlet-I, an indicator of entotic cell death^13^. Because we used Lysotracker staining to detect late-stage entotic structures throughout our study, we also incubated the cells with Lysotracker to confirm co-localisation of LAMP1 and lysosomal staining. As shown in Figure 3G, during internalisation, lysosomes in both inner and outer cells showed a synchronous movement for a short time, followed by simultaneous accumulation of both LAMP1 and Lysotracker staining. As entotic cell death progressed, LAMP1 and Lysotracker staining increased inside the nucleus of the inner cell and eventually co-localised with nuclear staining. As expected, quantification of LAMP1 and Lysotracker intensities verified a complete colocalisation at all stages (Figure 3I, SV6).

### Entosis induced by TRAIL requires the presence of Caspase-8

To investigate whether entosis induced by TRAIL required caspase-8 or activation of caspases, we performed large-scale HCS-based analysis and quantified the rate of apoptosis and entosis in the cells treated with or without TRAIL in the presence or absence of Y-27632 (ROCK 1/2 inhibitor), z-VAD-fmk (pan-caspase inhibitor) or z-IETD-fmk (‘caspase-8’ inhibitor). We found that Y-27632, alone or in combination with TRAIL, did not affect the rate of PI-positive cells (Figure 4A) or cells with cleaved IETD substrate (Figure S5A) indicating that inhibition of ROCK signalling pathway did not affect the induction of TRAIL-induced apoptosis in HCT116 cells. As expected, TRAIL treatment resulted in a significant increase in both PI-positive cells and cells with cleaved IETD substrate. Also, the presence of either z-VAD-fmk or z-IETD-fmk blocked TRAIL-induced increases in both PI-positive cells and cells with cleaved IETD substrate confirming that cell death was due to induction of apoptosis (Figure 4A, Figure S5A). Interestingly, as shown in Figure 4B, quantification of entotic structures showed that the presence of either z-VAD-fmk or z-IETD-fmk did not affect the rate of entosis in control or TRAIL-treated cells, suggesting that inhibition of caspase activity did not affect the rate of entosis. As expected^13^, co-treatment with Y-27632 significantly inhibited the rate of entosis induced by TRAIL-treatment. Taken together, these results indicate that apoptosis and entosis are two independent mechanisms both can be induced in response to TRAIL treatment.

**Figure 4.**
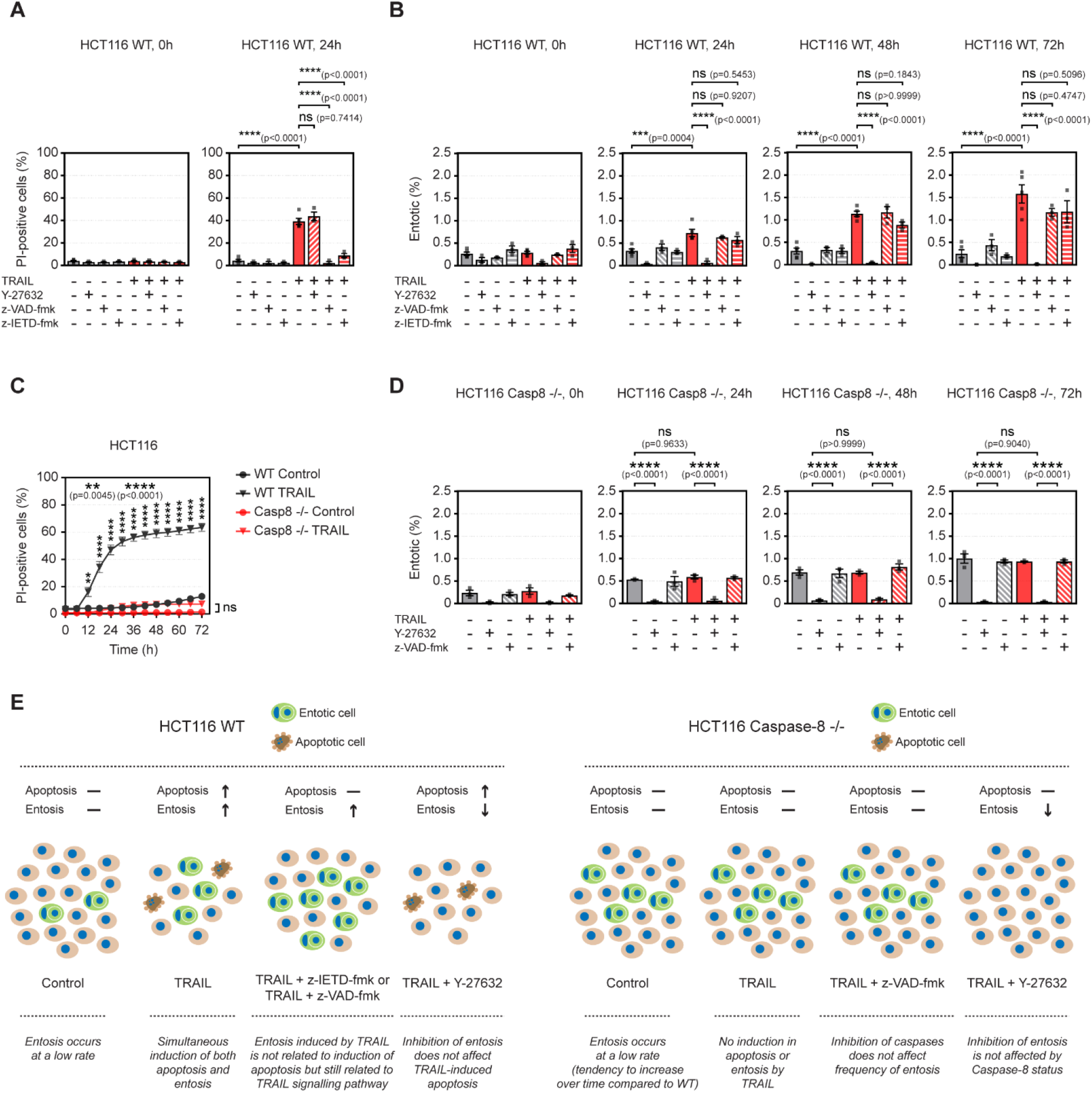
Entosis induced by TRAIL requires presence of Caspase-8. **A-B.** Apoptosis and entosis are independently induced by TRAIL treatment. **A.** Inhibition of entosis does not affect the rate of TRAIL-induced apoptosis. Quantification of PI-positive cells in HCT116 cells treated with or without TRAIL in the absence or presence of z-VAD-fmk, z-IETD-fmk or Y-27632 for 24 hours. Data are shown as individual values for each experiment as well as means ± S.E.M. from three experiments. Statistical significance was tested using one-way ANOVA followed by Tukey’s multiple comparison test, ****p<0.0001, ns: not significant. **B.** Inhibition of apoptosis does not affect the rate of TRAIL-induced entosis. Quantification of entotic structures in HCT116 cells treated with or without TRAIL in the absence or presence of z-VAD-fmk, z-IETD-fmk or Y-27632 for 72 hours. Data are shown as individual values for each experiment as well as means ± S.E.M. from at least three experiments. Statistical significance was tested using one-way ANOVA followed by Tukey’s multiple comparison test, ***p<0.001, ****p<0.0001, ns: not significant. **C-D.** Induction of both apoptosis and entosis by TRAIL requires presence of Caspase-8. **C.** Quantification of PI positive cells in HCT116 WT and Caspase-8 −/− cells treated with or without TRAIL for 72 hours. Data are shown as means ± S.E.M. from three experiments. **p<0.01, ****p<0.0001, ns: not significant. **D.** Quantification of entotic structures in Caspase-8 −/− cells treated with or without TRAIL in the absence or presence of z-VAD-fmk or Y-27632 for 72 hours. Data are shown as individual values for each experiment as well as means ± S.E.M. from three experiments. Statistical significance was tested using one-way ANOVA followed by Tukey’s multiple comparison test, ****p<0.0001, ns: not significant. **E.** Summary of changes in apoptosis and entosis rates in response to manipulation of TRAIL signalling pathway.

Caspase-8 sits at the nexus of death and survival pathways downstream of TRAIL signalling. In the past few years, a non-apoptotic (non-enzymatic) role of Caspase-8 has emerged in switching TRAIL signalling to survival pathway activation in cancer cells^10,11^. To examine whether Caspase-8 played a non-enzymatic role in the induction of entosis in TRAIL-treated cells, we quantified PI-positive cells and entotic structures in Caspase-8 CRISPR/Cas9 knockout cells^11^. As expected, TRAIL treatment did not change the rate of PI-positive cells in *CASP8* null cells as compared to controls (Figure 4C). Interestingly, we observed a trend for increased entosis events in control-treated *CASP8* null cells compared to control-treated WT cells over time, although the difference was significant only at 72 hours. Remarkably, we observed no difference in the rate of entosis between control- and TRAIL-treated *CASP8 null* cells indicating that Caspase-8 is required for TRAIL-induced entosis. We obtained similar results as in the wild type cells when we pre-treated *CASP8* null cells with z-VAD-fmk or Y-27632. z-VAD-fmk did not affect the entosis rate in either control- or TRAIL-treated cells, whereas Y-27632 significantly inhibited the rate of entosis in both (Figure 4D). Of note, when we cultured *CASP8 null* cells in SILAC RPMI (media with no glucose) for glucose deprivation (a known inducer of entosis^15^), we could still observe a significant increase in the rate of entosis indicating that involvement of caspase-8 in TRAIL-induced entosis is a specific mechanism that can be uncoupled from ‘canonical’ entosis signalling (Figure S5B). Overall, our results suggest that induction of entosis by TRAIL does not require the activation of downstream apoptotic signalling pathway and caspase-8 activation but rather requires the non-enzymatic presence of Caspase-8 (Figure 4E).

### Inhibition of apoptosis alters inner cell fate towards release

It has been reported that inhibition of apoptosis either genetically by overexpressing BCL-2^14^ or pharmacologically by using caspase inhibitors^23^ results in the release of inner cells. To examine this in the context of TRAIL treatment, we performed inner cell fate analysis using time-lapse microscopy (Figure 5A) after inhibiting apoptosis pharmacologically by z-VAD-fmk, or genetically by using *BAX/BAK* double knockout cells (Figure S6). We found that the majority of inner cells (70.34 ± 0.93 %) underwent entotic cell death in WT control cells, and TRAIL treatment did not affect the rate of inner cell death (65.26 ± 4.06 %). However, treatment with z-VAD-fmk, alone or in combination with TRAIL, resulted in a significant reduction in inner cell death and increased the rate of release in inner cells (Figure 5B, left). Strikingly, in *BAX/BAK* null cells, the vast majority of inner cells did not undergo entotic cell death, instead, they were released regardless of treatment (Figure 5B, right, SV7). There was no difference in inner cell division among treatments in WT and *BAX/BAK* null cells (Figure 5B). As expected, pretreatment with Y-27632 blocked the formation of entotic structures in controls as well as TRAIL-treated WT and *BAX/BAK* null cells (not shown). Collectively, our results indicate that inhibition of apoptosis signalling changed inner cell fate towards release.

**Figure 5.**
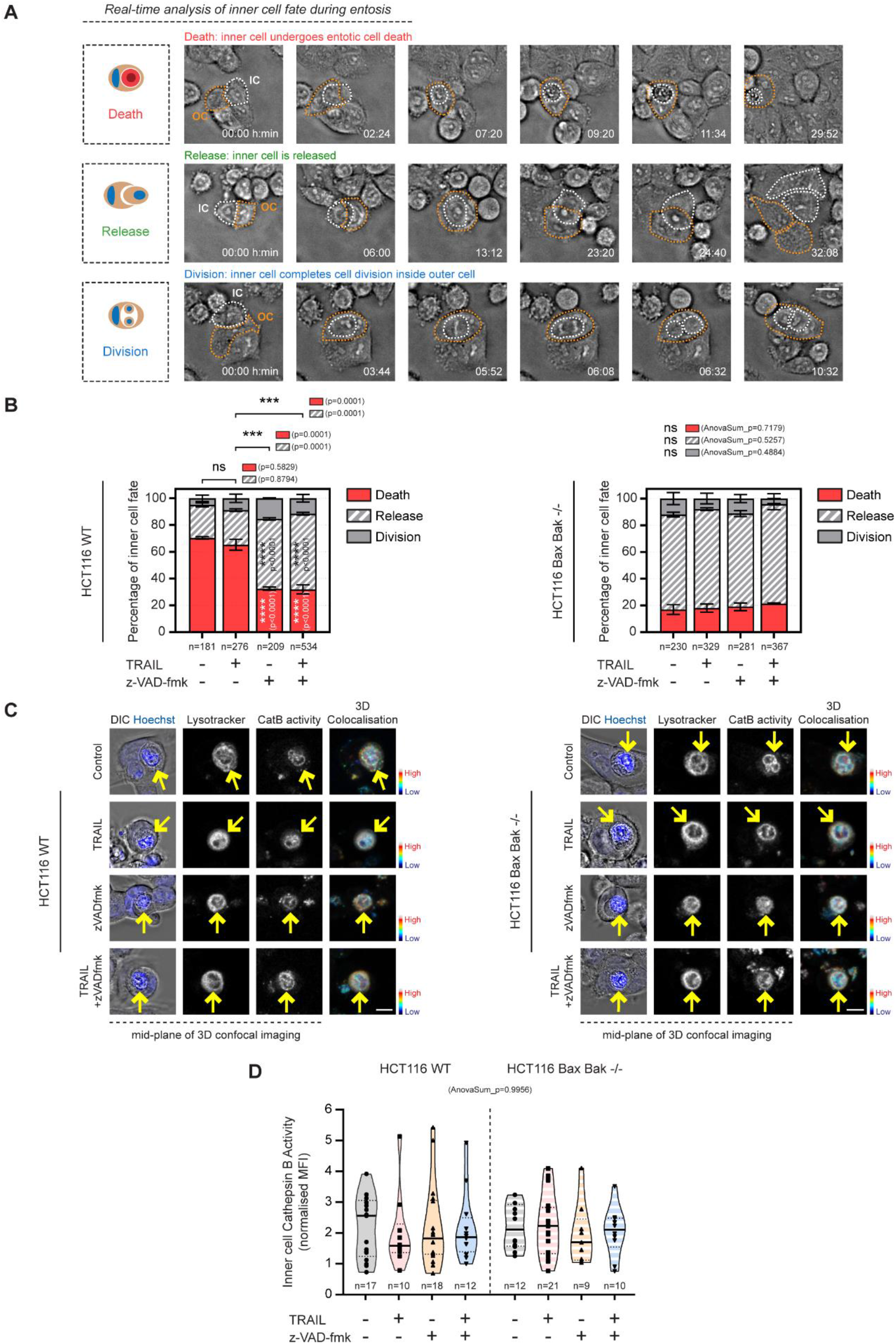
Deletion of Bax/Bak or Inhibition of caspase activation alters inner cell fate towards release. **A.** Schematic representation of inner cell fates and representative time-lapse images showing inner cells undergoing entotic cell death (Death), inner cell is released (Release) and inner cell completes cell division (Division). Inner cells (IC) and outer cells (OC) are indicated as orange-dashed and white-dashed lines, respectively. Images are representatives from three experiments, Scale bar: 10 μm. **B.** Quantification of inner cell fates in HCT116 WT and Bax Bak −/− cells treated with or without TRAIL in the absence or presence of z-VAD-fmk. Data are shown as individual values for each experiment as well as means ± S.E.M. n, total number of cells analysed over 48 hours from three experiments. Statistical significance was tested using one-way ANOVA followed by Tukey’s multiple comparison test. Asterisks inside individual bars indicate comparison*s* with control. ***p<0.001, ****p<0.0001, ns: not significant. **C.** Representative 3D confocal microscopy images of DIC, Hoechst (blue), lysotracker and fluorogenic Cathepsin B substrate of late stage entotic cells in WT and Bax Bak −/− cells treated with or without TRAIL in the presence or absence of z-VAD-fmk. 3D colocalization of lysotracker and Cathepsin B substrate is visualised. Yellow arrows indicate inner cells. Scale bars: 10 μm. Images are representatives from three experiments. **D.** Quantification of Cathepsin B activity in late stage entotic cells in WT and Bax Bak −/− cells treated with or without TRAIL in the presence or absence of z-VAD-fmk. Data are shown as individual values for each cell as well as median and quartiles, statistical significance was tested using unpaired two tailed t-test. n, total number of cells analysed from three experiments.

To further understand whether *Bax* and *Bak* are involved in regulating downstream entotic cell death, we examined Cathepsin B activity in late-stage entotic cells by 3D confocal microscopy. As shown in Figure 5C, inner cells exhibited Lysotracker accumulation co-localising with increased Cathepsin B activity in all treatments confirming that inner cells were undergoing entotic cell death during analysis. We observed no morphological difference in late-stage entotic cells among treatments in either WT or *BAX/BAK* null cells (Figure 5C). Likewise, quantification of the mean fluorescence intensity of Cathepsin B substrate cleavage yielded no significant change among treatments and cell lines (Figure 5D).

### Clinical association among TRAIL signalling, cell-in-cell structures and colorectal cancer

Next, we investigated the relationship between CICs, expression of proteins involved in TRAIL signalling, clinicopathological characteristics and outcome in a cohort of stage 2 and 3 CRC patients randomized to either observation or chemotherapy (5-FU/leucovorin) post-resection from the NI240 phase III clinical trial^24^. Demographic and clinicopathological characteristics of the CRC patient cohort are summarised in Supplementary File 1 page 19 (SF1, p19). We blindly quantified CICs in high-quality HE-stained TMA cores (Figure 6A, top) (n= 1410) from n=232 distinct patients prepared from the centre of the tumour (n=648), invasive front of the tumour (n=210) and adjacent normal tissue (n=552) (SF1, p21,22). Besides H&E staining, a combination of different cytoplasmic and nuclear staining has been used to detect CICs^17^. Therefore, we increased the sample size up to six consecutive TMA sections by quantifying CICs in the TMA cores (n=1381) of c-MET (stains both the cytoplasm and the cell membrane, Figure 6A, bottom) and nuclear staining (haematoxylin) prepared from the same tissues (centre of the tumour, n=636; invasive front of the tumour, n=210; matched normal, n=535). We observed a strong correlation when comparing cell-in-cell events identified from HE- and c-MET-stained TMA cores (Kendall rank correlation tau=0.46, p<0.0001, SF1, p11). Thus, for downstream analyses, we pooled CICs estimated from either staining and considered them as biological replicates, totalling up to 2 and 6 cores per patient for tissue from the invasive front and tumour centre/matched normal tissue, respectively (SF1, p10, 21, 22). Notably, the distribution of CICs in tumours showed a high heterogeneity among different sections within the same patients as well as across patients (Figure 6B).

**Figure 6.**
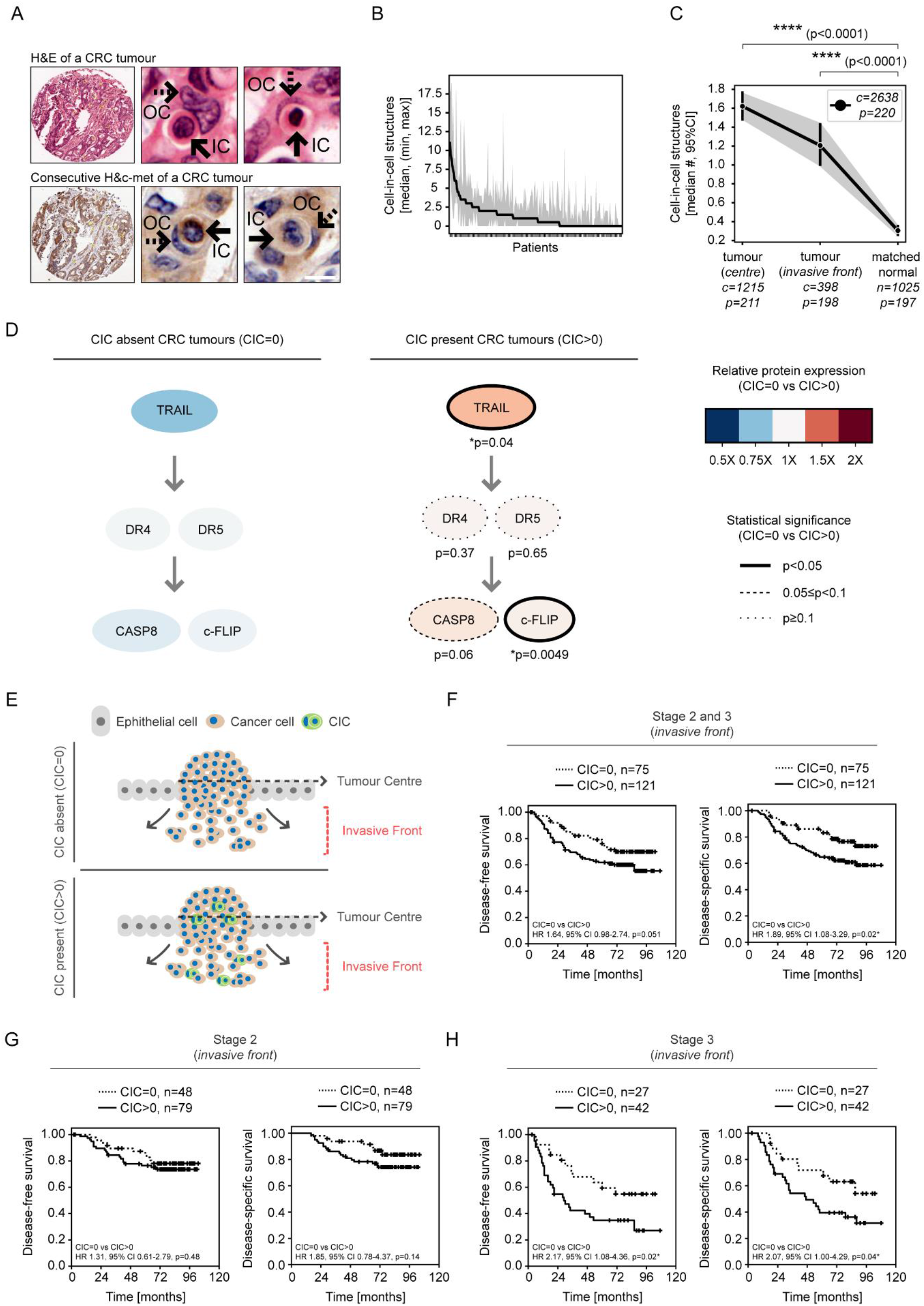
Clinical association among TRAIL signalling, CICs and colorectal cancer. **A-C.** CICs are increased in CRC tumours compared to matched normal tissues. **A.** Consecutive H&E (top) and H&c-MET (bottom) sections showing representative CICs in a CRC tumour. Arrows indicate inner (plain arrows) and outer cells (dashed arrows), Scale bar: 10 μm. **B.** Distribution of CICs in TMA sections of CRC patients (n=254). **C.** Quantification of CICs in TMA sections prepared from tumour centre, invasive front and matched normal tissues. *c* and *p* indicate number of tumour cores and patients, respectively. ****p<0.0001. **D.** Overexpression of proteins involved in TRAIL signalling is associated with presence of CICs in CRC tumours. Association between expression levels of TRAIL, DR4, DR5, CASP8, c-FLIP, and CICs. Relative protein expression between CIC absent (median CIC=0) and CIC present (median CIC>0) tumours is indicated in colour. Statistically significant differences in protein expression between CIC absent and CIC present patients are encoded by the Node-Edge style graphs (circles with solid line, dashed line and dotted line indicate p<0.05, 0.05≤p<0.1 and p≥0.1, respectively). Individual p values are shown under each protein. *p<0.05. **E-H.** Presence of CICs in the invasive front region of tumours is an independent prognostic marker for disease-free survival and disease-specific survival in colorectal cancer. **E.** Schematic representation of the centre and the invasive front of a tumour, and absence or presence of CICs. **F-G.** Kaplan-Meier estimates of Disease-free survival and Disease-specific survival of the NI240 cohort (F) comparing stage 2 (G) and 3 (H) CRC patients grouped by the presence or absence of CICs in the invasive front region. *p<0.05.

We then focussed on patients with CIC quantifications from both tumour centre, invasive front and matched normal tissues (n=188 patients). We found that the number of cell-in-cell events differed by location (p<0.0001) decreasing from the tumour centre to the invasive front (β −0.06, 95% CI-0.23 to 0.12, ns) and further into the matched normal tissue (β −2.36, 95% CI −2.78 to –1.94), (Figure 6C; SF1, p24). We further explored the relationship between CICs by tissue and clinicopathological features. We found no statistically significant difference in between CICs and TNM staging, age, sex, lymphovascular invasion and treatment (SF1, p25). However, we observed fewer CICs in rectal tumours compared to those from the colon (β −0.47, 95%CI –0.82 to −0.12, p=0.009; SF1, p13, p15, p25).

Next, we examined the association between CICs and proteins involved in TRAIL signalling pathway including TRAIL, TRAIL/R1 (DR4), TRAIL/R2 (DR5), c-FLIP and Caspase-8 in tumour tissue (Figure 6D). In agreement with other reports^17,19^ a proportion of patients (75 of 211, 36%) showed no evidence of CICs in tumour tissues. Thus, we used the absence or presence of CICs as cut-off and classified the CRC patients in our cohort as CIC absent (CIC=0) or CIC present (CIC>0) if the median number of CICs detected across multiple biological replicas within the same tissue was equal to or greater than 0, respectively. Remarkably, we found that CIC-positive tumours had higher expression of TRAIL (CIC>0 (ref. CIC=0): 0.59, 95% CI 0.02 to 1.15, p=0.04) and c-FLIP (CIC>0 (ref. CIC=0): 0.78, 95% CI 0.00 to 1.55, p=0.049), with a trend towards higher expression of CASP8 (CIC>0 (ref. CIC=0): 0.66, 95% CI −0.02 to 1.35, p=0.06).

We next investigated the association between CICs and clinical outcome with Kaplan-Meier estimates and (un)adjusted Cox regression hazard models. We found no statistically significant difference in disease-free (DFS) and disease-specific (DSS) (we dropped OS from plots) survival curves when comparing patients with CICs (CIC>0) and those with no CICs (CIC=0) detected in tumour centre when examining the whole cohort or when stratifying by stage (log-rank p>0.05; SF1, p15-p19). Intriguingly however, patients with CICs (CIC>0) in the invasive front of the tumours had an approximately two-fold increase risk of relapse (HR 1.64, 95% CI 0.98 to 2.74, p=0.051) and cancer-related death (HR 1.89, 95% CI 1.08 to 3.29, p=0.02) compared to patients with no detected CICs (CIC=0) (Figure 6G). Moreover, the presence/absence of CICs in the invasive front of the tumour was an independent prognostic marker after controlling for baseline clinicopathological characteristics (DFS: HR 2.01, 95% CI 1.19 to 3.39, p=0.007; DSS: HR 2.13, 95% CI 1.21 to 3.74, p=0.006; SF1, p27). We then assessed whether the association between CICs and clinical outcome differed by stage (2 vs 3). We found that the prognostic value of CICs in the invasive front region was seen only in stage III patients (log-rank p-values=0.03 and 0.04 for DFS and DSS, respectively) (Figure 6H, I). In a multivariate analysis controlling for baseline clinicopathological characteristics, among stage III patients, those CIC-positive showed an increased risk of relapse (HR 2.67, 95% CI 1.31 to 5.43, p=0.004) and cancer-related mortality (HR 2.33, 95% CI 1.12 to 4.87, p=0.02), (SF1, p28). These associations were null among stage II patients. Taken together, these results indicate that the invasive front of tumours is the region where CICs have clinical relevance in stage III CRC patients.

## Discussion

Recent studies have revealed that well-documented inducers of apoptosis can simultaneously activate multiple cell death and survival mechanisms in cancer cells, such as autophagy^25^, necroptosis^26^ and entosis^18^. In this study, we have discovered that colorectal cancer cells can simultaneously initiate entosis and apoptosis upon TRAIL stimulation. The TRAIL signalling pathway is activated by immune effector cells, such as CTLs and NK cells and is also a promising therapeutic target for cancer treatment; however, frequently only a proportion of cancer cells die by apoptosis despite exposure to saturating concentrations of TRAIL^8^. Cell-to-cell variability, stochastic genetic fluctuations and non-genetic differences among individual cells, seems to be the driving factor resulting in the fractional killing. Here, measurement of single-cell caspase activation kinetics and MOMP upon TRAIL stimulation in HCT116 cells corroborated previous findings^20^ that TRAIL treatment results in heterogeneous apoptosis activation. Strikingly, while analysing the time of MOMP in single cells, we discovered that, instead of showing a typical sharp decrease during apoptosis^27^, a small proportion of cells displayed a dramatic increase in TMRM signal. Although TMRM has been widely used in staining mitochondrial membrane potential in living cells^20^, we could not find any study in the literature reporting a similar behaviour of TMRM. This unanticipated behaviour led us to track these cells in time-lapse images and revealed that TMRM accumulation occurred after a cell invading into another cell, a process known as entosis. Co-staining of TMRM and lysosomes demonstrated that TMRM accumulation was coincident with the acidification of inner cells during entotic cell death. Intriguingly, just before the accumulation due to entotic cell death, TMRM exhibited a punctate formation around inner cells.

In this study, we also improved some of the current limitations of entosis quantification by designing a large-scale HCS-based approach, which allows simultaneous detection of apoptosis and entosis continuously without requiring any processing step such as washing or trypsinisation. Our fully customisable approach randomly takes several fields of view and enables quantification of a large number of cells (~50.000 cells/treatment) over time. It automatically detects the late-stage entosis events, and most importantly, cells are monitored in their natural state in culture conditions throughout the experiment. Entosis was initially found to be induced by matrix detachment in human mammary epithelial cells and breast cancer cells^13^. Further studies have revealed that several factors can induce entosis such as myosin activation^28^, epithelial cadherin expression^15^, glucose starvation^15^, radiation, and chemotherapeutics such as paclitaxel, oxaliplatin and cisplatin^18^ in attached cell lines. Our HCS-based quantification demonstrated that TRAIL stimulation promoted entosis in colorectal cancer cells in a time-dependent manner. We were able to confirm entosis induction by TRAIL also in different cell line models as well as spheroids. Intriguingly, the presence of CHX significantly increased TRAIL-induced apoptosis but reduced the rate of entosis. Given that CHX has been shown to reduce key anti-apoptotic proteins such as Mcl-1 and c-FLIP^29^, induction of entosis by TRAIL might be related to pro-survival signalling. Pre-treatment with the pan-caspase inhibitor z-VAD-fmk inhibited TRAIL-induced caspase activation and apoptosis, however, it did not affect the induction of entosis by TRAIL. On the contrary, pre-treatment with the ROCK inhibitor Y-27632 inhibited TRAIL-induced entosis but did not affect caspase activation or induction of apoptosis by TRAIL. Thus, our results suggest that, although entosis and apoptosis can co-occur in a population of cells, they seem to be two independent mechanisms.

While apoptotic cells show caspase activation and DNA fragmentation, entotic cell death involves recruitment of autophagic lipidation machinery proteins including LC3, ATG5, ATG7 and VPS34 onto entotic vacuole, followed by recruitment of LAMP1 leading to lysosomal fusion ultimately resulting in inner cell death by cathepsin B release^14^. Inner cells however also showed features of apoptosis signalling during TRAIL treatment such as caspase activation, cytochrome c release and membrane blebbing. Strikingly, when entotic cell death started taking place, apoptotic features were not detectable after a certain stage, possibly due to acidification of inner cells. Markers such as cleaved caspase-3 and cytochrome c formed foci when lysosomal accumulation occurred during entotic cell death. Florey *et al.* showed that inner cells can also upregulate autophagy to survive within outer cells^14^, and it has been well-documented that inner cells can divide inside outer cells or can be released^30^. Although activation of apoptosis does not seem to be involved in the induction of entosis in many systems, accumulating evidence suggests that blocking apoptosis either pharmacologically by caspase inhibitors^23^ or genetically by overexpressing anti-apoptotic genes such as BCL-2^14^ results in the release of inner cells. Similarly, in this study, inner cell fate analysis showed that the vast majority of inner cells underwent entotic cell death unless co-treatment with z-VAD-fmk or knock out of *BAX* and *BAK* altered inner cell fate towards survival and release. Our experiments suggest that, despite the fact that lysosomal cell death will eventually kill a proportion of inner cells during TRAIL-induced entosis, the presence of *BAX* and *BAK*, as well as caspase activity may still be necessary for entotic cell death to occur. Carrying a cell inside may have a huge impact on outer cells, resulting in a population-level survival response. The presence of inner cells can lead to cytokinesis failure in outer cells and result in aneuploidy^31^. Moreover, entotic cell death provides a survival advantage to outer cells under stress condition such as glucose starvation^15^. In agreement with these studies, we observed that the majority of outer cells survived TRAIL treatment, i.e. that cells showing limited caspase activation may become outer cells in response to TRAIL receptor activation.

Following TRAIL stimulation, activation of procaspase-8 at the DISC requires its homodimerisation. Recent studies have revealed that caspase-8 plays a non-enzymatic role in activating pro-survival TRAIL signalling leading to the induction of NF-κB-dependent cell survival and release of several cytokines including IL-8, CXCL5, CXCL1 and CXCL2 from the surviving cells. Release of these cytokines has been shown to create a tumour-supportive microenvironment and promote tumour growth^10^. We showed that pre-treatment with z-IETD-fmk, a selective caspase-8 inhibitor, inhibited TRAIL-induced caspase activation and apoptosis, however, did not affect the rate of entosis induced by TRAIL. Remarkably, CRISPR/Cas9-mediated deletion of *CASP8* resulted in a reduction in TRAIL-induced entosis as well as apoptosis in HCT116 cells, providing another biological example for a non-enzymatic role of caspase-8.

Despite CICs having been reported in a variety of human cancers for almost a century^12^, understanding their clinical relevance has progressed very little because studies have been mostly occasional reports showing examples in the tumour tissues. Until now, to the best of our knowledge, there have been no studies assessing whether tumours have an increase in CICs compared to normal tissues in the same patients. Here, in addition to addressing this question, we performed a comprehensive clinical analysis of CICs in colorectal cancer patient samples. In line with the literature^19^, the number of CICs greatly varied among consecutive TMA sections from the same patients as well as among patients, suggesting that multiple sections should be used to get a robust quantification of CICs in clinical samples. Intriguingly, colorectal cancer tumours showed significantly more CICs compared to matched normal tissues. Notably, CICs were not associated with clinicopathological features of CRC including TNM staging, age, sex, lymphovascular invasion, or treatment. Remarkably, overexpression of TRAIL and c-FLIP but not DR4 and DR5 were associated with the presence of CIC in CRC tumours. These tumours also showed a trend to express higher levels of caspase-8. Taken together, in addition to our *in vitro* findings, our results suggest that TRAIL signalling is also clinically associated with the presence of CICs in CRC patients.

CICs have been reported not only in the centre of tumours but also in the invasive front region in cancers^19^. However, whether the presence of CICs in different intratumoural regions has prognostic value was unknown. We show that the prognostic value of CICs emerges from their presence in the invasive front region of tumours but not in the tumour centre in colorectal cancer. The invasive front region is where the most progressed cells of a tumour invade into the immune cell-rich stromal region^32^. Due to higher immune cell content, this region may possess higher cytokine (and death ligands) content and might reflect greater interactions between immune cells and cancer cells ^33^. CRC cells have shown to escape from NK-cell mediated tumour surveillance by acquiring resistance to TRAIL-mediated apoptosis *in vitro* (HCT116) and *in vivo*^34^. The *in vitro* as well as clinical data presented in this study might be a motivation for further investigations to evaluate whether CICs play a role in this immune cell – tumour cell interaction in the invasive front region. The invasive front region is also especially important in CRC because it is the region where tumour budding, an established prognostic factor for CRC patients, is observed^35^. Because tumour buds consist of isolated tumour cells or small tumour cell clusters that detach (lose adhesion) from the central tumour and migrate into the stroma^36^, it would be interesting to examine whether entosis plays a role in the formation of these structures. Our analysis also revealed that the prognostic value of CICs in colorectal cancer arises from the strong association in stage 3 patients. We did not detect this association in stage 2 CRC patients who however showed a lower recurrence rate^37^. Because CICs show a potential to be used as an independent prognostic marker, there is an urgent need to investigate the biological and the clinical relevance of CICs in the invasive front region in other solid tumours.

## Methods

### Reagents, antibodies and plasmids

Dimethyl sulfoxide (#D8418), Dulbecco’s Phosphate Buffered Saline (#D8537), Fetal Bovine Serum (FBS, #F7525), G-418 (#04727878001), Hanks’ Balanced Salt solution (#H8264), Hoechst 33258 (#94403), L-Glutamine (#G7513), Penicillin-Streptomycin (#P0781), Propidium iodide (#P4864), RPMI-1640 Medium (#R0883), Trypsin-EDTA (#T4049), Z-Arg-Arg-AMC (#C5429) were purchased from Sigma Aldrich (Darmstadt, Germany). Lipofectamine 2000 (#11668019), Lysotracker Deep Red (#L12492), Lysotracker Red (#L7528), Tetramethylrhodamine − Methyl Ester (#T668), DRAQ5 (#65-0880-96), Alexa Fluor 488 and 647 secondary antibodies (#A11008, #A11029, #A21202, #A21206, #A31571, #A31573), β-Catenin antibody (#RB-9035) were obtained from Thermo Fisher Scientific (Dublin, Ireland). Tumour necrosis factor-related apoptosis-inducing ligand (TRAIL) (#310-04) was from PeproTech (London, England). SiR-Hoechst (#SC007) was purchased from Tebu-bio (Peterborough, England). Y-27632 (#S1049), z-VAD-fmk (#S7023) and z-IETD-fmk (#S7314) were from Selleck Chemicals (Munich, Germany). LC3A/B (#4108) and Cleaved Caspase-3 (#9661) antibodies were purchased from Cell Signaling Technology (Leiden, The Netherlands). Cytochrome c (#556432) antibody was from BD Biosciences (Oxford, England). EM Goat anti-Rabbit IgG (#EM.GAR10) was purchased from BBI Solutions (Crumlin, UK). Lamp1-mScarlet-I was a gift from Dorus Gadella (Addgene plasmid #98827). pSCAT8 (CFP-IETD-Venus) was previously generated by our group^20^. Concentrations of drugs and inhibitors used in the experiments are as follows unless otherwise mentioned, TRAIL:100 ng/ml, CHX: 1-2 μg/ml, z-IETD-fmk: 25 μM, z-VAD-fmk: 25, 50 μM, Y-27632: 20 μM.

### Cell culture and transfections

HCT116 cells were cultured in RPMI-1640 supplemented with 10% FBS, 2 mM L-Glutamine, 100 units/ml penicillin and 100 μg/ml streptomycin at 37°C in a humidified atmosphere at 5% CO_2_. Cells were cultured in 6-well plates until they reached ~70% confluency, then transfected with 0.6 μg of plasmid DNA (CFP-IETD-Venus) and 6 μl of Lipofectamine 2000 in 1 ml Opti-MEM at 37°C with 5% CO_2_ for 4-12 hours. To generate stable cell lines, transfected cells were cultured in the presence of 0.5 − 1 mg/ml G-418 for 1-2 weeks, and fluorescent clones were picked and expanded. For the generation of stable HCT116 Lamp1-mScarlet-I cell line, cells were transfected with 0.75 μg of plasmid DNA (Lamp1-mScarlet-I) and 7.5 μl of Lipofectamine 2000 in 1 ml Opti-MEM. HCT116 Bax Bak −/−^38^ cells and their wild types were a kind gift from Richard J. Youle (National Institutes of Health, Bethesda, MD, USA), HCT116 Caspase-8 CRISPR/Cas9 knockout cells and their wild types^11^ were a generous gift from Seamus Martin (Trinity College Dublin, Dublin, Ireland). HCT116 cells have been authenticated before the beginning of the study and after completion of the experiments, and regularly checked for mycoplasma.

### Live-cell time-lapse microscopy and FRET measurements

Cells were seeded on sterile 12-mm glass-bottom WillCo-dishes (WillCo Wells B.V., Amsterdam, Netherlands) and grown in RPMI medium at 37 °C with 5% CO_2_ for 24 hours until they attached to the glass surface. Cells were then incubated in staining medium containing a combination of 1 μg/ml Hoechst 33258 or 1 nM SiR-Hoechst or 1 μM DRAQ5, 30 nM TMRM, 50 nM Lysotracker Green, Lysotracker Red or Lysotracker Deep Red depending on the experiment. WillCo-dishes containing stained cells were covered with embryo tested sterile mineral oil and mounted on an LSM 710 Confocal laser-scanning microscope (Carl Zeiss Ltd, Cambridge, UK) equipped with a 40×/1.3 numerical aperture Plan-Apochromat oil immersion objective and a microscope incubator chamber (37 °C with 5% CO_2_) (Pecon, Erbach, Germany) or Nikon TE2000, Automated Epi-fluorescence Microscope. UV or cyan fluorescence was excited using 405 nm (detection range 415-494 nm or 450-505nm, respectively), green or yellow fluorescence was excited using 488 nm (detection range 490-544 nm or 505-560nm, respectively), red fluorescence was excited using 561 nm (detection range 590-644 nm) and far-red fluorescence was excited using 633 nm (detection range 638-728 nm) lasers, respectively. Images were recorded at a resolution of 1024×1024 pixels for 16 to 72 hours with 4 to 20 minutes intervals. Treatments were added after ~30 minutes of the baseline measurement. Cell fates were determined from time-lapse microscopy images of DIC and Hoechst recorded for 48 hours. Image processing (background subtraction, median filtering, creating masks, ratiometric calculations) was carried out using Fiji/ImageJ (version 1.52i, Wayne Rasband, NIH, Bethesda, MD, USA). Kinetics of single-cell IETD substrate cleavage were analysed by combining time-lapse ratiometric imaging data generated by dividing CFP by FRET after background subtraction.

### High Content Screening

#### Imaging

Cells were seeded on 96-Well optical-bottom plates (#165305, Thermo Fisher Scientific, Dublin, Ireland) at a density of 7500 cells/well and grown in RPMI medium at 37 °C with 5% CO_2_ for 24 hours until they attached to the surface, then stained with Hoechst (1 μg/ml), Propidium Iodide (PI) (1 μg/ml) and Lysotracker Deep Red (50 nM). The dyes were present in the medium throughout the experiment. Ninety-six well plates containing stained cells were treated in triplicates as indicated in figure legends and imaged using Cellomics ArrayScan VTi (Thermofisher, Surrey, UK) equipped with a 10x PlanApo objective lens (NA 0.45), multi diode light source (Lumencor SOLA SE II, AHF, Germany) set to 20-22% of maximal emission intensity and a monochrome CCD camera (Hamamatsu Orca AG). Time series of images (12 fields of view per well) were collected from each well for 72 hours with 6 hours interval, at a resolution of 1024×1024. Dye concentrations and image acquisition rate were optimized in advance to minimize phototoxicity. Hoechst was imaged using 100 ms exposure time, a 381-394 nm excitation filter and a 415-460 nm emission filter; CFP was imaged using 1000 ms exposure time, a 422-432 nm excitation filter and a 457-487 nm emission filter; FRET was imaged using 300 ms exposure time, a 422-432 nm excitation filter and a 528-555 nm emission filter; PI was imaged using 75 ms exposure time, a 582-596 nm excitation filter and a 608-683 nm emission filter; Lysotracker Deep Red was imaged using 500 ms exposure time, a 644-657 nm excitation filter and a 690-730 nm emission filter. The autofocus module on Cellomics ArrayScan was applied on the FRET channel in every third field of view at 4×4 binning.

#### Image processing and analysis

All images were segmented and measured using CellProfiler 2.2 (Broad Institute, Cambridge, Massachusetts, USA). We optimized and combined modules for uneven/non-uniform background illumination correction, nuclei isolation and shape/intensity parameter identification to create an HCS image processing and analysis workflow. The data obtained was further processed using Excel 2016 (Microsoft, California, USA). The multiplexing of CFP-IETD-Venus, Hoechst, PI and Lysotracker Deep Red staining allows the automated identification of specific cell states. These include cells, which exhibit caspase-8-like activity (FRET disruption caused by IETD cleavage), cells showing an apoptotic nuclear morphology (PI negative small and fragmented nuclei), dead cells (PI positive), late entotic cells (bright Lysotracker Deep Red stain that co-localized with Hoechst stain). Because we were not able to automatically identify early entotic cells, they were manually quantified based on detecting round-shaped Hoechst stained cells inside a large vacuole within another Hoechst stained cell showing a crescent-shaped nuclear morphology as shown in Fig 3A.

### Immunofluorescence staining and 3D confocal imaging

Cells were seeded on sterile 12-mm glass-bottom WillCo-dishes (WillCo Wells B.V., Amsterdam, Netherlands) and grown in RPMI medium at 37 °C with 5% CO2 for 24 hours until they attached to the glass surface. Cells were then treated for indicated times and fixed in 4% paraformaldehyde diluted in PBS for 15 min at room temperature. For experiments shown in Fig 3 (A and B), Lysotracker Deep Red was added 30 minutes before fixation. After washing three times in 1X PBS for 5 minutes each, cells were blocked in a blocking buffer (1X PBS, 5% Normal Goat Serum and 0.3% Triton X-100) for 60 minutes. Primary antibodies were prepared in a dilution buffer (1X PBS, 1% BSA, 0.3% Triton X-100) and samples were incubated overnight at 4 °C. For double immunofluorescence staining, a mixture of two primary antibodies from different species were simultaneously added. After washing three times in 1X PBS for 5 minutes each, complementary secondary antibodies (different fluorochromes were used for double immunofluorescence staining) prepared in a dilution buffer and cells were incubated for 2 hours at room temperature in the dark. Samples were washed in 1X PBS for 5 minutes each, stained with 2 μg/ml for 10 minutes and washed once with 1X PBS. Samples were then mounted on an LSM 710 confocal laser-scanning microscope (Carl Zeiss Ltd, Cambridge, UK) fitted with a 63×/1.4 NA Plan-Apochromat oil immersion objective. 405 nm, 488 nm, 561 nm and 633 nm lasers were used to excite DAPI, Alexa Fluor 488, Alexa Fluor 561 and Alexa Fluor 647 conjugated secondary antibodies with detection ranges of 415-494 nm, 490-544 nm, 560-628 nm and 638-728 nm respectively. Z-stacks were taken by recording a series of optical slices with an optimal thickness (0.6 - 1 μm) at 1024×1024 or 2048×2048 pixels throughout the entire cells. Further image processing (background subtraction, median filtering and construction of orthogonal views) and quantification were carried out using Fiji/ImageJ (version 1.52i, Wayne Rasband, NIH, Bethesda, MD, USA). For experiments shown in Fig 1I, mean intensity values for outer cells were normalised to mean intensity values of apoptotic cells recorded from the same field of view. For experiments shown in Fig 3C, mean intensity values for inner cells were normalised to the average of five neighbouring cells from the same field of view.

### Cathepsin B activity

Cells were seeded on sterile 12-mm glass-bottom WillCo-dishes and grown in RPMI medium at 37 °C with 5% CO2 for 24 hours until they attached to the glass surface. After treatments, Z-Arg-Arg-AMC, Fluorogenic substrate for cathepsin B, was added to the media at a concentration of 50 μM. Cells were fixed in 4% paraformaldehyde or imaged alive on LSM 710 Confocal laser-scanning microscope (Carl Zeiss Ltd, Cambridge, UK) as described above.

### Correlative Light and Electron Microscopy (CLEM) analysis

Cells were seeded on 35 mm sterile high Grid-500 μ-Dishes (ibidi GmbH, Munich, Germany), grown in RPMI medium at 37 °C with 5% CO2 for 24 hours until they attached to the surface, then treated with TRAIL for 48 hours. After staining with Lysotracker for 30 minutes, cells were immediately fixed in 4% paraformaldehyde and stained with Hoechst. 3D confocal imaging of DIC, Hoechst and Lysotracker staining were performed as described above and areas containing entosis events were recorded for further analysis. Samples were then fixed again in 2.5% glutaraldehyde in sodium cacodylate buffer then washed three times in cacodylate buffer. After fixation, samples were placed in 1% osmium in sodium cacodylate buffer for 60 minutes then rinsed three times in distilled water. Next, samples were incubated in increasing concentrations of methanol (50%, 60%, 70%, 80%, 90%, 95%, 100%, 100%). Subsections of interest (SOI) were isolated from the samples using a scalpel and placed in LR white resin prepared in methanol (1:1 ratio) for 60 minutes, followed by LR white resin only for another 60 minutes. SOI was then embedded in fresh LR white resin using a gelatine capsule and incubated overnight under UV light at 4°C. Ultramicrotome sections at 60-90 nm thickness were generated using a Leica EM UC6 (Laboratory Instruments and Supplies, Ashbourne, Ireland) and trapped on a pioloform coated 3 mm 200 thin bar copper grids. Images of the events of interest were recorded using a Hitachi H-7650 transmission electron microscope (Hitachi, Tokyo, Japan) at an accelerating voltage of 40-100 kV. Enhance Local Contrast (CLAHE) plugin in Fiji/ImageJ (version 1.52i, Wayne Rasband, NIH, Bethesda, MD, USA) was used to automatically enhance the contrast of images shown in Fig3D-F.

### Analysis of CICs in colorectal cancer tissue microarrays

#### Patients cohort

Patients were drawn down from the NI240 phase III clinical trial coordinated at Queen’s University (Belfast, UK) where primarily stage 2 and 3 colorectal cancer patients were randomized to observation or adjuvant treatment with 5-FU/leucovorin following primary tumour resection^24^. Clinico-pathological characteristics including treatment randomization, TNM staging, tumour location, lymphovascular invasion and outcome were annotated for n=240 patients who met the inclusion criteria (SF1, p10). Patients were followed-up post-surgery for approximately 7 years (SF1, p20).

#### Tissue microarrays

Tissues from the tumour centre, the invasive front and matched normal mucosa were collected during surgical resection and preserved in formalin-fixed paraffin-embedded (FFPE) blocks for the construction of tissue microarrays (TMAs)^37^. Multiple TMA cores per patient (2 cores for the invasive front and up to 6 cores for normal and tumour centre) were stained for haematoxylin/eosin (HE) and Hematoxylin/Tyrosine-Protein Kinase Met (c-MET), totalling n=2791 cores in n=232 patients (SF1, p21). CICs that exhibited evidence of a complete internalisation of an entire cell were manually quantified by independently detecting at least four of the five following criteria: nucleus of inner cell (1), cytoplasm of inner cell (2), a crescent-shaped nucleus of outer cell (3), cytoplasm of outer cell (4), visible vacuolar space between inner and outer cells (5). Events identified as CICs were recorded using QuPath v0.2.0-m2^39^. Moreover, multiple TMAs cores (n=7601 from tumour centre and matched normal tissue were stained for TRAIL, TRAIL-R1 (DR4), TRAIL-R2 (DR5), Caspase-8, and c-FLIP for n=236 patients (SF1, p22). Protein expression was computed as previously described ^37^ as the product of the staining scores for intensity (0-3 integer scale) and coverage (0-4 integer scale), with low and high values indicating weak and strong expression, respectively.

In downstream analysis, we included n=220 patients with annotated clinical records and CICs in at least a tissue type. Additional analysis examining the association between CICs and protein profiles was possible in n=220 patients with materials to quantify the proteins (SF1, p15). Data and analysis code are publicly available and archived at Zenodo at https://doi.org/10.5281/zenodo.3841833.

### Statistical analysis

#### In vitro data

When comparing groups containing single-cell information, data obtained from three experiments were presented as individual values for each cell as well as median and quartiles in each group (Fig 1D, Fig 1H, Fig 1I, Fig 1K, Fig 3C) by Violin Plots. When comparing groups containing a value per experiment (Fig 2B and 2C, Fig4 A-D), data were presented as individual values for each experiment as well as the standard error of the mean value (SEM) in each group by Aligned Dot Plots with Bars. Statistical significance between the means of two groups was tested using unpaired two-tailed t-test, and among the means of more than two groups was tested using one-way ANOVA followed by Tukey’s multiple comparison test. GraphPad Prism 8 Software (La Jolla, California, USA) was used to perform statistical comparison tests and preparation of graphs. Figures were prepared using Adobe Illustrator Software (version CS6, San Jose, California, United States). Asterisks represent significance (*p<0.05; **p<0.01; ***p<0.001, ****p<0.0001), “ns” indicates non-significance. All experiments were repeated at least three times unless otherwise indicated.

#### Clinical data

##### CIC events analysis

Agreement between cell-in-cell events (median across biological replicas per tissue per patient) estimated from HE or c-MET staining was assessed by Kendall rank correlation tau (python package scipy, version 1.2.0). We analysed the association between CIC events by tissue and clinical covariates with zero-inflated Poisson regression models, including a random effect for each patient and covariate-fixed effects. We reported effect sizes (estimate), 95% confidence intervals (CIs) and p-values computed by likelihood ratio tests (R package glmmTMB, version 0.2.3).

##### Protein expression analysis

Protein expression across biological replicas per tissue and per patient was aggregated by the median. Association between protein expression and cell-in-cell events (binary, CIC=0 vs. CIC>0) was assessed in tumour tissue by unadjusted linear models.

##### Outcome analysis

We examined disease-free (DFS) and disease-specific (DSS)survival as outcome end-points. DFS, DSS and OS were computed as the time interval between surgical resection and either study end-date or end-point event (tumour recurrence as evidenced by radiological imaging or cancer-related death, respectively). Patients with no events were censored at study end-date. Differences in survival curves between patient groups were assessed by Kaplan-Meier estimates and logrank tests. We investigated the association between CICs [absence (CIC=0) vs. presence (CIC>0)] with the outcome by univariate and multivariate Cox regression hazard models. The multivariate model was adjusted by baseline patient characteristics selected a priori, including stage (2 vs 3), treatment (observation vs. 5-FU/leucovorin), age (continuous), sex (male vs. female), tumour location (colon vs. rectum). We reported hazard ratios (HRs), 95% CIs and p-values computed by loglikelihood ratio tests. Survival analyses were performed in python with the lifelines package (version 0.23.6)^40^.

Asterisks represent significance (*p<0.05; **p<0.01; ***p<0.001, ****p<0.0001), “ns” indicates non-significance between groups in the figures. p-values were not adjusted for multiple comparisons as these analyses were deemed exploratory.

## Acknowledgements

This study was supported by Science Foundation Ireland (14/IA/2582; 16/RI/3740; 16/US/3301) and the Health Research Board (16/US/330; TRA/2007/26). We wish to sincerely thank the NI240 study participants and trial investigators.

## Author Contributions

Conceptualisation: EB, HD, JHMP, Methodology: EB, HD, BLC, MS, Formal Analysis: EB, HD, MS, Investigation: EB, HD, MS, Resources: JHMP, SVS, DBL, SM, Writing - original Draft: EB; Writing – Review&Editing: HD, MS, SVS, DBL, SM, JHMP, Visualisation: EB, MS. Supervision: JHMP, Funding Acquisition: DBL and JHMP.

