## Supplementary Figures for "TRAIL signalling promotes entosis in colorectal cancer"

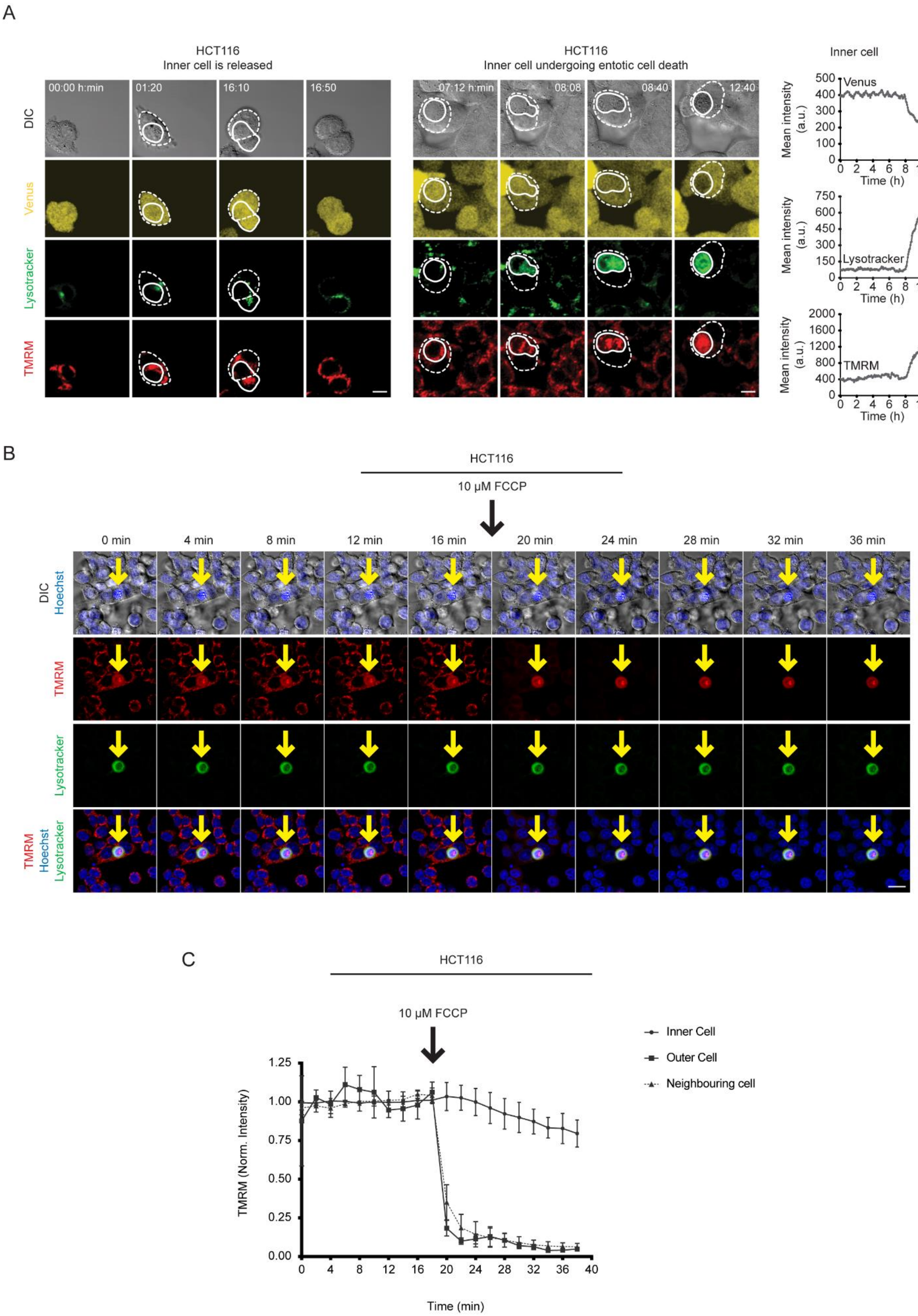
A
HCT116
Inner cell is released
00:00 h:min
01:20
16:10
16:50
DIC
Venus
Lysotracker
TMRM
HCT116
Inner cell undergoing entotic cell death
07:12 h:min
08:08
08:40
12:40
Inner cell
Venus
Lysotracker
TMRM
Mean intensity (a.u.)
Time (h)
B
HCT116
10 μM FCCP
0 min
4 min
8 min
12 min
16 min
20 min
24 min
28 min
32 min
36 min
DIC Hoechst
TMRM
Lysotracker
TMRM Hoechst Lysotracker
C
HCT116
10 μM FCCP
Inner Cell
Outer Cell
Neighbouring cell
TMRM (Norm. Intensity)
Time (min)

**Supplementary Figure 1.**

**A.** Representative time-lapse images of DIC, Venus, Lysotracker and TMRM showing an inner cell being released (left) and inner cell undergoing entotic cell death (right). Accumulation of TMRM and Lysotracker coincides with the loss of Venus signal in the entotic inner cell (right). Straight and dashed lines indicate inner and outer cells, respectively. Images are representatives from 3 experiments. Scale bar: 20 µm.
**B.** Representative time-lapse images of DIC, Hoechst, TMRM and Lysotracker showing the effects of FCCP treatment on inner cell TMRM signal during entotic cell death. **C.** Quantification of TMRM mean fluorescence intensity in the corresponding inner, outer and neighbouring cells. Images are representatives from 3 experiments. Yellow arrows indicate inner cell. Scale bar: 20 µm.

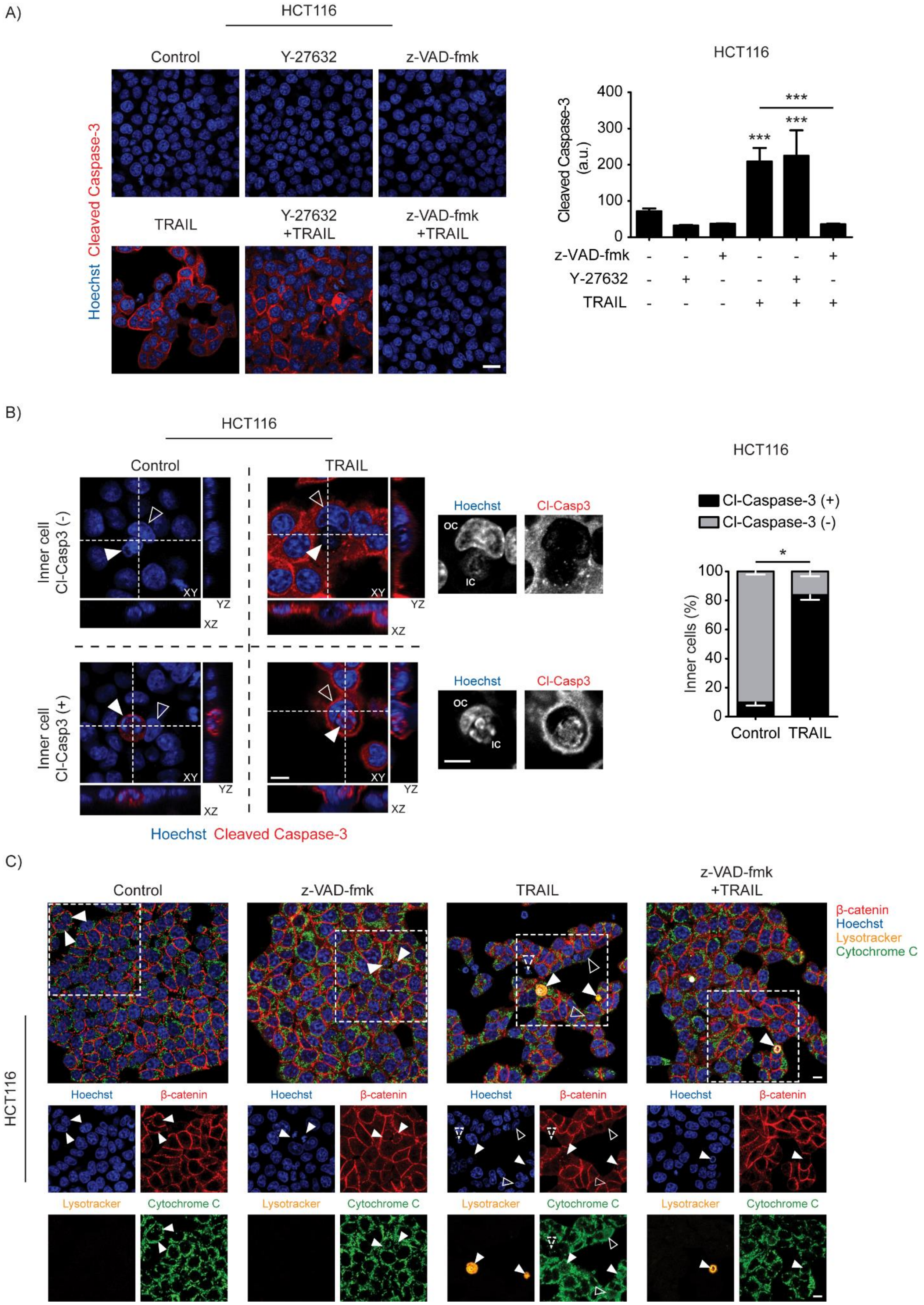
A)
HCT116
Control
Y-27632
z-VAD-fmk
TRAIL
Y-27632 +TRAIL
z-VAD-fmk +TRAIL
Hoechst Cleaved Caspase-3
HCT116
Cleaved Caspase-3 (a.u.)
400
300
200
100
0
***
z-VAD-fmk
Y-27632
TRAIL
B)
HCT116
Control
TRAIL
Inner cell Cl-Casp3 (-)
Inner cell Cl-Casp3 (+)
XY
YZ
XZ
Hoechst
Cl-Casp3
OC
IC
Hoechst Cleaved Caspase-3
HCT116
Cl-Caspase-3 (+)
Cl-Caspase-3 (-)
*
Inner cells (%)
100
80
60
40
20
0
Control
TRAIL
C)
Control
z-VAD-fmk
TRAIL
z-VAD-fmk +TRAIL
β-catenin
Hoechst
Lysotracker
Cytochrome C
HCT116

**Supplementary Figure 2.**

**A.** Representative confocal microscopy images of Hoechst (blue) and Cleaved Caspase-3 (red) staining in HCT116 cells treated with or without TRAIL in the absence or presence of z-VAD-fmk and Y-27632. **B.** Quantification of Cleaved caspase-3 mean fluorescence intensity in HCT116 cells treated with or without TRAIL in the absence or presence of z-VAD-fmk and Y-27632. Statistical significance was tested using one-way ANOVA followed by Tukey's multiple comparison test (n=3). ***p<0.001. Scale bar: 10 µm.

**B.** Representative 3D confocal microscopy images of Hoechst (blue) and Cleaved Caspase-3 (red) staining showing Cleaved Caspase-3 positive and negative inner cells in control and TRAIL-treated cells (left). Quantification of inner cell Cleaved Caspase-3 status in control and TRAIL-treated cells. White and black arrowheads indicate inner and outer cells, respectively. Statistical significance was tested using unpaired two tailed t-test (n=3). *p<0.05. Scale bar: 20 µm.

**C**. Representative confocal microscopy images of β-catenin (red), Hoechst (blue), Lysotracker (orange) and Cytochrome C (green) staining in HCT116 cells treated with or without TRAIL in the absence or presence of z-VAD-fmk. White, black and dashed arrowheads indicate inner cells, cells showing Cytochrome c release and apoptotic cells, respectively. Images are representatives from 3 experiments. Scale bar: 10 µm.

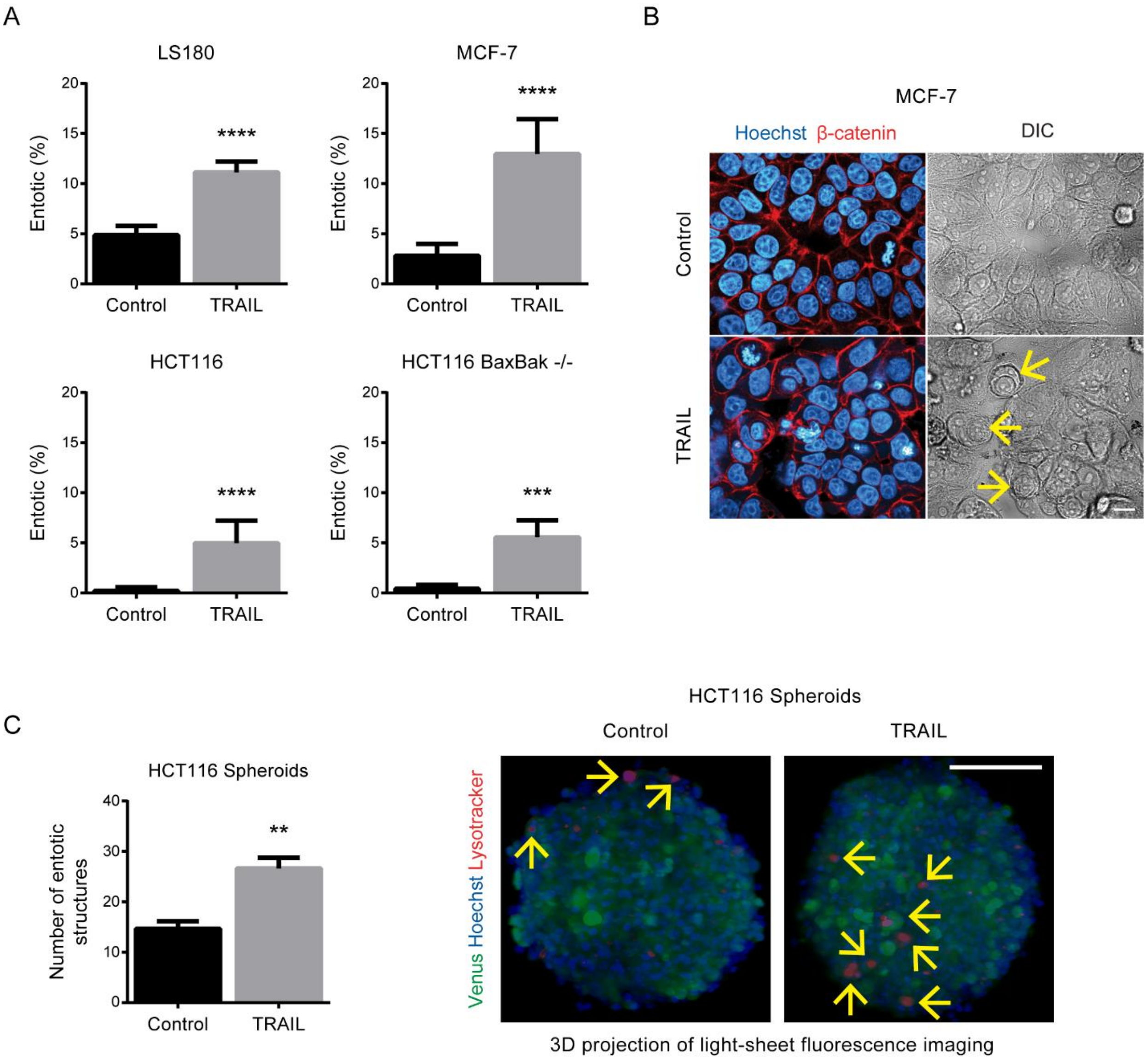


**Supplementary Figure 3.**

**A.** Quantification of entotic events in LS180, MCF7, HCT116 and HCT116 Bax Bak -/- cells treated with control or TRAIL for 48 hours. **B.** Representative confocal microscopy images of DIC, Hoechst (blue) and β-catenin (red) staining in MCF-7 cells treated with control or TRAIL. Yellow arrows indicate entosis events. >500 cells were quantified from at least two experiments. Statistical significance was tested using unpaired two tailed t-test, **p<0.01, ***p<0.001, ****p<0.0001. Scale bar: 20 µm.

**C.** Quantification of entotic events in HCT116 spheroids treated with control or TRAIL (left). **B.** 3D projections of light sheet fluorescence imaging of Venus (green), Hoechst (blue) and Lysotracker (red) in control and TRAIL-treated HCT116 spheroids. Three spheroids per treatment were quantified. Yellow arrows indicate lysotracker positive inner cells. Statistical significance was tested using unpaired two tailed t-test, **p<0.01. Scale bar: 100 µm.

**A**

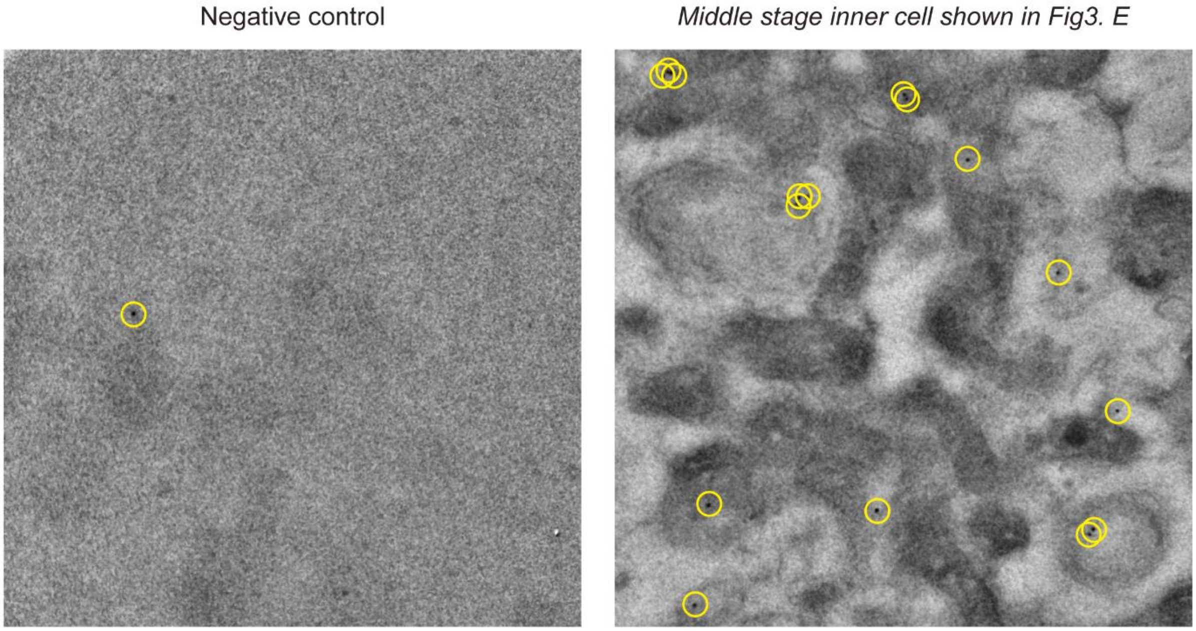


**B**

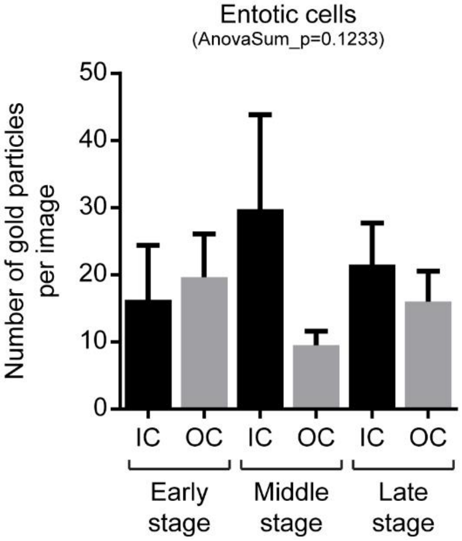


**Supplementary Figure 4.**

**A.** Representative TEM images showing ultrastructural localisation of LC3 in negative control and middle stage inner cell. Yellow circles indicate gold particles. **B.** Quantification of gold particles in early, middle and late stage inner (IC) and outer cells (OC) shown in Figure 3. Statistical significance was tested using one-way ANOVA followed by Tukey's multiple comparison test.

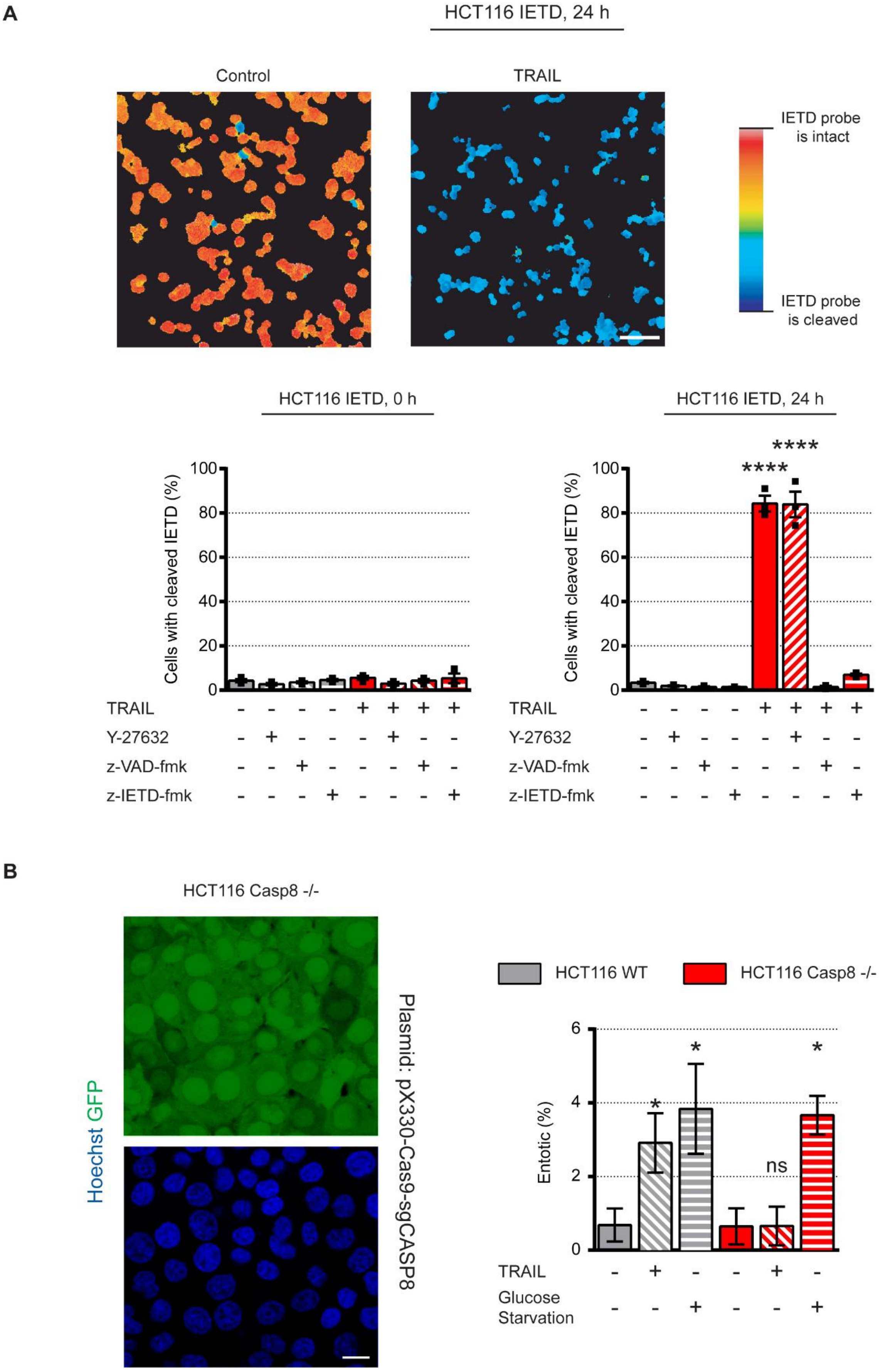
A
HCT116 IETD, 24 h
Control
TRAIL
IETD probe is intact
IETD probe is cleaved
HCT116 IETD, 0 h
HCT116 IETD, 24 h
Cells with cleaved IETD (%)
****
****
TRAIL
Y-27632
z-VAD-fmk
z-IETD-fmk
B
HCT116 Casp8 -/-
Hoechst GFP
Plasmid: pX330-Cas9-sgCASP8
HCT116 WT
HCT116 Casp8 -/-
Entotic (%)
ns
TRAIL
Glucose Starvation

**Supplementary Figure 5.**

**A.** Representative images (top) and quantification of cells with cleaved IETD substrate (bottom) in HCT116 cells treated with or without TRAIL in the absence or presence of z-VAD-fmk, z-IETD-fmk or Y-27632. Statistical significance was tested using one-way ANOVA followed by Tukey's multiple comparison test (n=3). ****$p<0.0001$. Scale bar: 20 µm.

**B**. Representative images of Hoechst-stained (blue) HCT116 Caspase-8 CRISPR -/- cells (green) (left). Quantification of entotic events in controls, TRAIL-treated or glucose-starved HCT116 WT and Caspase-8 CRISPR -/- cells (right). >1000 cells were quantified from two experiments. Statistical significance was tested using one-way ANOVA followed by Tukey's multiple comparison test. *$p<0.05$, ns: not significant. Scale bar: 20 µm.

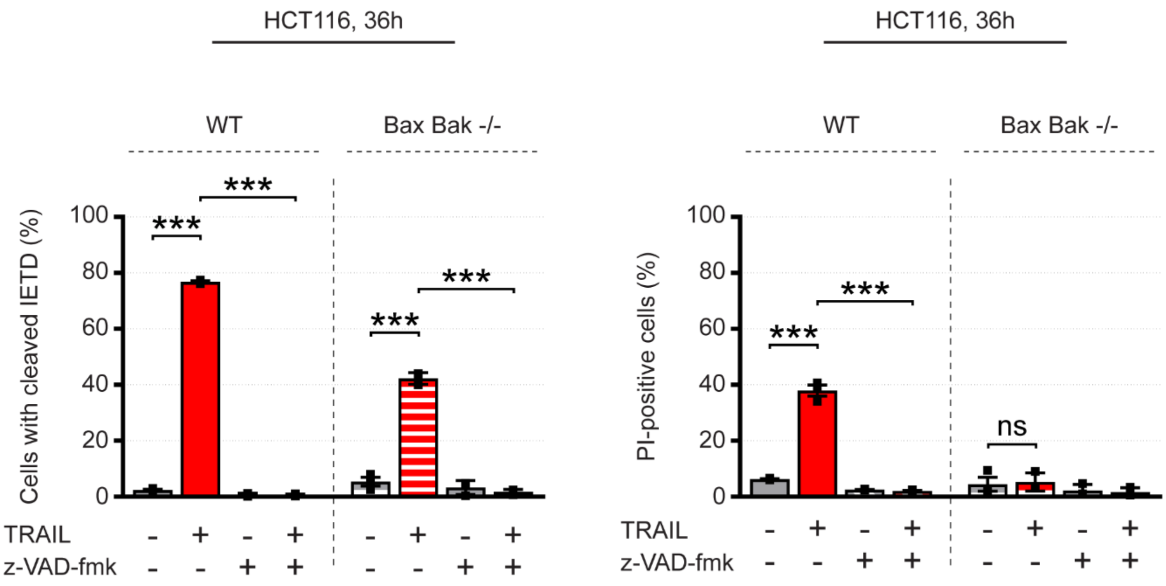


**Supplementary Figure 6.**

Quantification of Cells with cleaved IETD substrate and PI-positive cells in HCT116 Bax Bak -/- treated with or without TRAIL in the absence or presence of z-VAD-fmk. Statistical significance was tested using one-way ANOVA followed by Tukey's multiple comparison test (n=3), ***p<0.001, ns: not significant.

### Supplementary Methods

#### Entosis quantification

Cells were seeded, stained and treated on sterile 12-mm glass-bottom WillCo-dishes (WillCo Wells B.V., Amsterdam, Netherlands) dishes in RPMI medium at 37 ºC with 5% $CO_2$. Entosis events were determined by counting at least 500 cells/sample by either using time-lapse microscopy images or 3D confocal microscopy of immunofluorescence staining (Hoechst and β-catenin) after 48 hours of treatment. Events were quantified based on detecting round-shaped (Hoechst stained) cells inside a large vacuole within another (Hoechst stained) cell showing a crescent-shaped nuclear morphology as shown in Fig 3A. Both dead and alive inner cells were quantified as entotic.

#### Light Sheet Fluorescence Microscopy

***Sample preparation***: HCT116 cells were seeded on 96-Well spheroid plates (96-well Black/Clear Round Bottom Ultra-Low Attachment Spheroid Microplate, Corning, UK) at a density of 1000 cells/well and grown in RPMI medium at 37 ºC with 5% $CO_2$ for 48 hours until they formed spheres. To stain the spheres homogenously stains were added to the spheroid culture as indicated for at least 5hrs at the following concentrations: Lysotracker Red 50 nM, Lysotracker Deep Red 10 nM, sirHoechst 0.5 µM, Hoechst 33258 1µg/ml.

***Imaging:*** The spheroids were embedded in 1% low melting agarose in PBS (Sigma Aldrich, Ireland) at 38-40 °C, sucked into a glass capillary while liquid. When hardened the capillary was mounted in the microscope sample holder and chamber of the Light Sheet Fluorescence Microscope (Lightsheet Z1, Carl Zeiss, Germany) using glass capillaries with an inner diameter of 1.0 mm and outer diameter of 1.5 mm. A plunger was then used to push the agar with the embedded spheroids out of the capillary into the liquid in front of the 20x 1.0 NA lens. The light sheet was generated with two 10 x 0.2 NA lenses illuminating the sample alternating from each side using the pivot scan mode. Images were taken at zoom 1.0 to 1.3 using a light sheet thickness from 3.61 µm to 4.11 µm respectively (405 nm excitation). Hoechst 33258 was excited using the 405 nm laser line, the Venus component of the IETD FRET probe was excited using 488 nm, Lysotracker Red with 561 nm, sirHoechst with 638 nm, all using the 405/488/561/640 nm notch filter in the emission light path to block scattered laser light. To split the emission onto the two PCO edge sCMOS cameras filter cubes with beam splitter one at 490 nm using band pass filters of 420 to 470 nm (Hoechst/CFP) and 505-545 nm (FRET) and beam splitter two at 510 nm using a band pass filters of 575-615 nm (Lystotracker Red) and beam splitter three at 560 nm using a band pass filter of 505 to 545 nm (Venus) and a long pass filter of 660 nm (sirHoechst). Image stacks were then taken moving the object along the optical axis of the imaging objective at 1µm steps and subsequently turning the object in 45 ° steps and re-aligning the object for each of the next 4 stacks. All images were processed using ZEN black and FiJi (ImageJ 1.52r-t). The multi view fusion and deconvolution were performed using the dedicated FiJi plugins (MultiView Reconstruction v5.0.20[1]) with either using Hoechst stained nuclei or Lysotracker stained lysosomes as alignment aid for registration. For the image fusion processing the scale was set to 2x using a server with dual Xeon Gold and 192 GB RAM (Power Edge R 740 XD, Dell EMC, Ireland), deconvolution was done with a $512^3$ pixel block size. 3D projections are presented in the figures.

**Immunogold Labelling**

Following TEM analysis, grids were incubated in a drop of blocking buffer (0.1 % Tween20, 2% Goat Serum, 2% BSA in PBS) for 30 mins at room temperature. After blocking, grids were placed in a drop containing LC3 primary antibody (1:100) for overnight at 4 °C. Next, grids were washed in a drop of PBS for an hour (4 x 15 mins). Following washing, grids were incubated in a drop of secondary antibody conjugated to 10-nm gold particles (1:50) for 1 hr at room temperature. Grids were washed in PBS for 25 mins (5 x 5 mins) and immediately imaged using a Hitachi H-7650 transmission electron microscope. Antibodies used in immunogold labelling were diluted in blocking buffer.
