## Supplementary Videos for "TRAIL signalling promotes entosis in colorectal cancer": SV_Legends.docx

Supplementary Video 1: HCT116 IETD cells showing formation of apoptotic and entotic structures in response to TRAIL treatment. TMRM and Venus are pseudocoloured as red and green. Arrows indicate entotic structures. Corresponding to Figure1E-G.

Supplementary Video 2: HCT116 cells showing surviving and dying entotic cells in response to TRAIL treatment. Arrows indicate entotic structures. Merge of DIC, Hoechst (blue) and Lysotracker (red) is shown. Corresponding to Figure1J.

Supplementary Video 3: 3D projection of an inner cell showing Lystoracker (red) accumulation during entosis. Hoechst and Mitotracker are pseudocoloured as blue and green. Corresponding to Figure1.

Supplementary Video 4: Reduction of Venus (yellow) signal coincides with accumulation of TMRM (red) and Lysotracker (green) in inner cells during entotic cell death. Corresponding to Supplementary Figure1.

Supplementary Video 5: 3D projection of HCT116 IETD spheroids showing entotic structures in control and TRAIL treatment. Venus (green), Hoechst (blue) and Lytostracker (red) are presented. Lysotracker stained nuclei indicate late stage entotic cells. Corresponding to Supplementary Figure3C.

Supplementary Video 6: Lysosomal events during entotic cell death in TRAIL-treated cells. HCT116 cells transfected with LAMP1-mscarlet-I (red) and stained with Hoechst (blue) and Lysotracker (green). Corresponding to Figure3G.

Supplementary Video 7: Release of inner cells in HCT116 Bax Bak -/- during TRAIL treatment. Arrows indicate inner cells. Corresponding to Figure5.
