## Supplementary File for "TRAIL signalling promotes entosis in colorectal cancer"

### TRAIL signalling promotes entosis in colorectal cancer - Supplementary File 1

2020-08-13

**References** **29**

### Abbreviations

- **cMET**: hematoxylin/tyrosine-protein kinase met staining;
- **CASP8**: caspase 8 protein;
- **CIC**: cell-in-cell events;
- **CI(s)**: confidence interval(s);
- **CRC**: colorectal cancer;
- **DFS**: disease-free survival;
- **DR4**: death receptor 4;
- **DR5**: death receptor 5;
- **DSS**: disease-specific survival;
- **FLIP**: FLICE (FADD-like IL-1B-converting enzyme)-inhibitory protein;
- **HE**: hematoxylin/eosin staining;
- **HR**: hazard ratio;
- **IHC**: immunohistochemistry;
- **NI240**: Northern Ireland phase III clinical trial;
- **TMA**: tissue microarray;
- **TRAIL**: TNF-related apoptosis-inducing ligand.

### Software packages

Complete datasets and analysis code are available via a public BitBucket repository and archived with Zenodo at 10.5281/zenodo.3841833. Analysis was performed in python (Van Rossum and Drake 2009), matlab (MATLAB 2014) and R (R Core Team 2020). The full list of packages and their versions are listed in the repository binder folder. Key libraries, corresponding programming language and usage in this study are listed in the table below.

| Language | Description | Package |
|---|---|---|
| Python | Data ingestion, cleaning and wrangling | *pandas* (McKinney and others 2010), *numpy* (Oliphant 2006) |
| Python | Data visualization | *matplotlib* (Hunter 2007), *seaborn* (Waskom et al. 2018), *upsetplot* (Alexander Lex 2014), *pydot* (wrapper for *graphviz* (Ellson et al. 2001)), *svgutils* |
| Python | Statistical analysis | *scipy* (Virtanen et al. 2020), *statsmodels* (Seabold and Perktold 2010), *tableone* (Pollard et al. 2018), *lifelines* (Davidson-Pilon 2019) |
| R | Statistical analysis | *glmmTMB* (Brooks et al. 2017), *car* (Fox and Weisberg 2019) |
| Matlab | Data visualization | *HCP (HeatmapCovariatePlot)* (Salvucci and Prehn 2019) |

### Figure panels for Figure 6 in the main manuscript

#### Figure 6 panel B

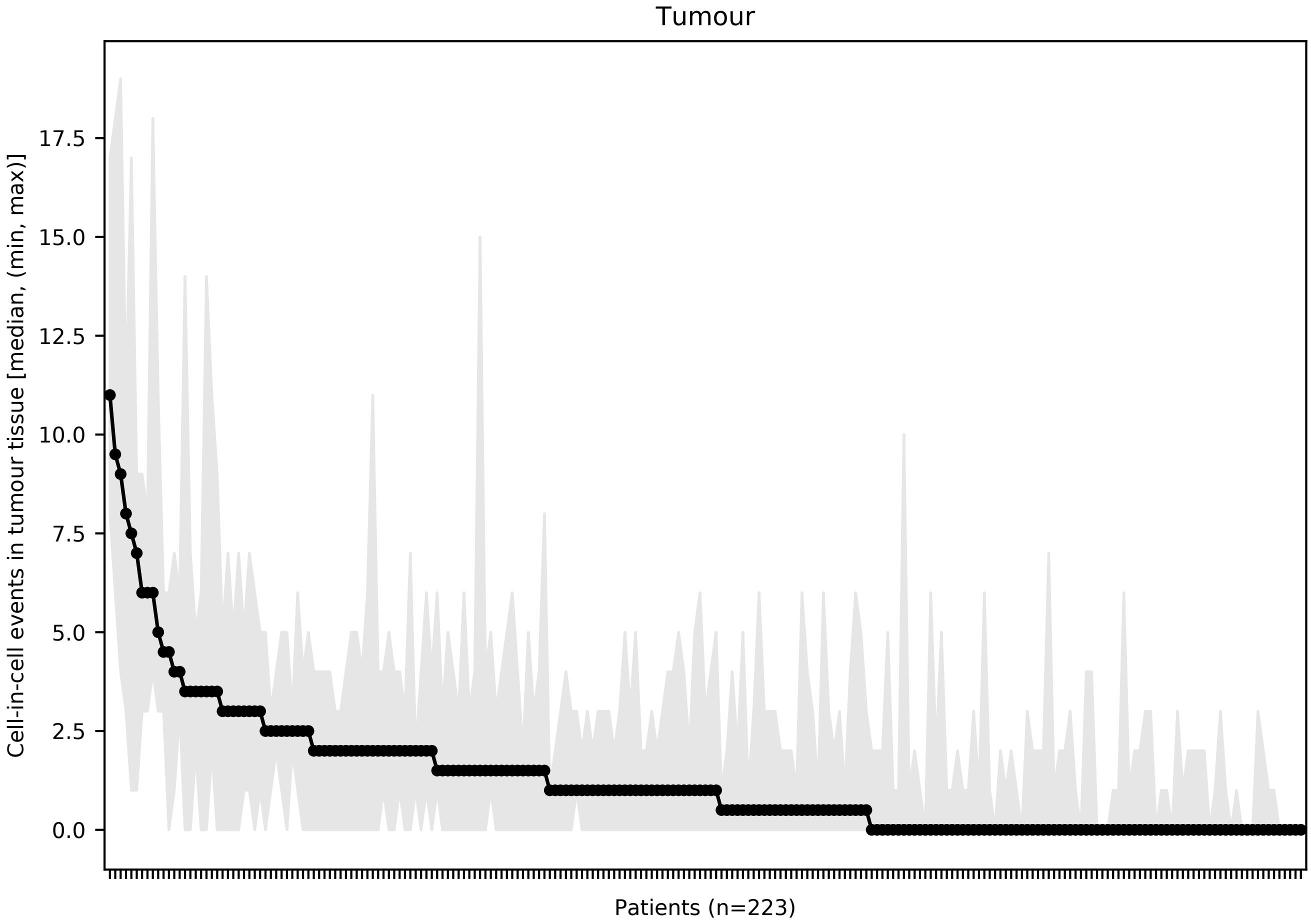

**Fig. 6B.** Inter- and intra-patient heterogeneity in cell-in-cell (CIC) events detected in tumor tissue and computed for each patient of the NI240 cohort. Patients (x-axis) are sorted in decreasing order of median CIC events (y-axis) detected in individual TMA cores prepared from tumour tissue and stained with either HE or cMET. Marker and shaded area indicate the median and the minimum/maximum CIC across the cores for each patient, respectively.

#### Figure 6 panel C

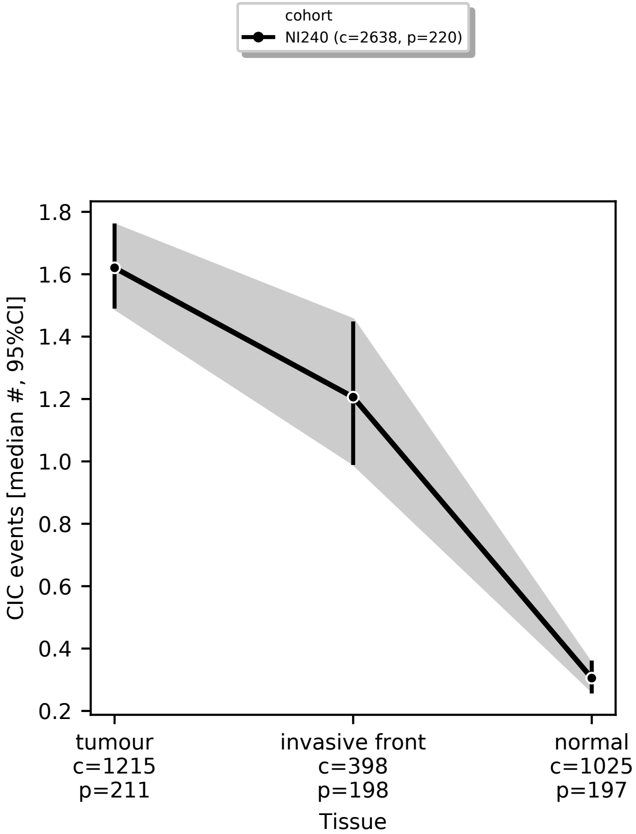

**Fig. 6C.** CIC events (median and 95% CI) detected in TMA sections prepared from tumour centre, invasive front and matched normal tissue. The letters "c" and "p" are abbreviations for "cores" and "patients", respectively. Corresponding statistical analysis is reported in **Sup. Table 6**.

**Figure 6 panel D**

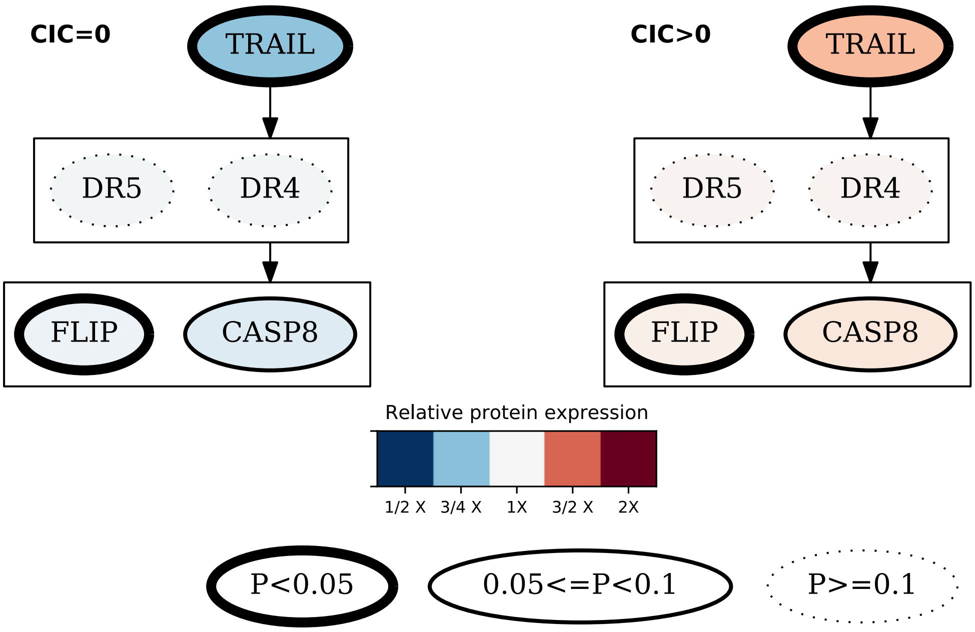

**Fig. 6D.** Association between expression of proteins involved in TRAIL signalling (TRAIL, DR4, DR5, CASP8 and FLIP) and CIC events. Relative protein expression between patients classified as CIC negative (CIC=0) or CIC-positive (CIC>0) based on the median CIC events observed across TMA sections prepared from tumour tissue is indicated in color. Red and blue shades indicate increased expression in CIC-positive or CIC-negative patients, respectively. Statistical significant differences in protein expression between CIC-negative and CIC-positive patients are encoded by the node edge style where solid tick lines and dotted thin lines indicate significant and non-significant differences, respectively. Visualization was generated with the python package *pydot*, built on *graphviz* (Ellson et al. 2001)). Complete statistical results are reported in **Sup. Table 8**.

#### Figure 6 panel F

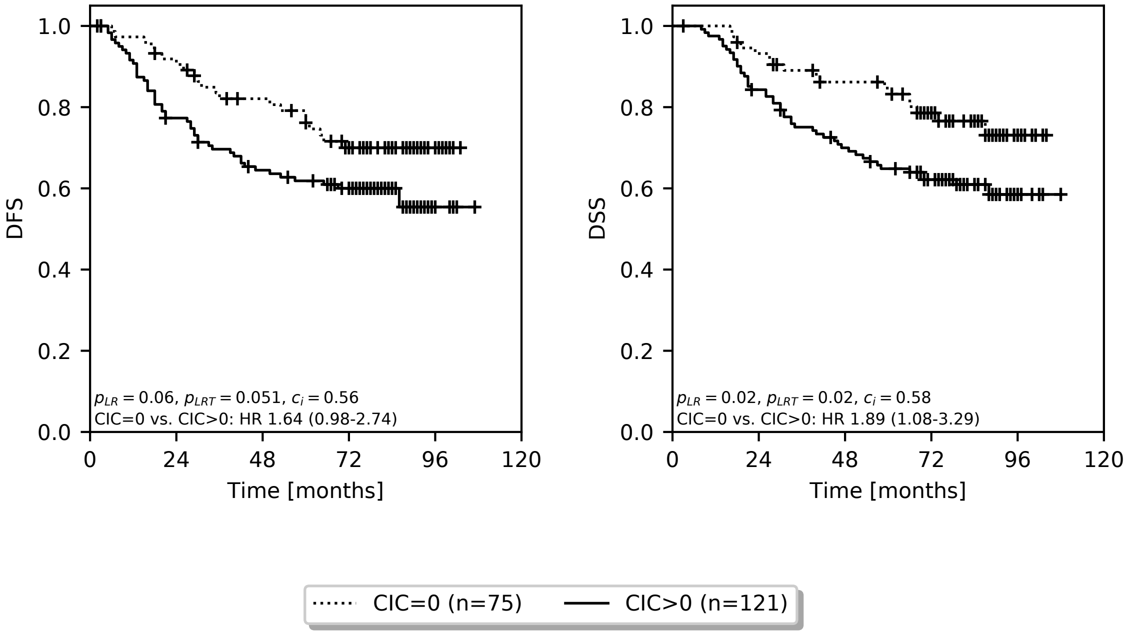

**Fig. 6F.** Kaplan-Meier estimates of DFS and DSS comparing stage II and III patients of the NI240 cohort grouped by the absence or presence of CIC events detected in the invasive front tissue. Patients were classified as CIC-negative (CIC=0) or CIC-positive (CIC>0) if the median number of CIC events detected across multiple biological replicas (TMA sections) prepared from the invasive front tissue was equal to or greater than zero, respectively. Kaplan-Meier estimators, logrank p-values and univariate Cox regression models were computed with the python package *lifelines*.

#### Figure 6 panel G

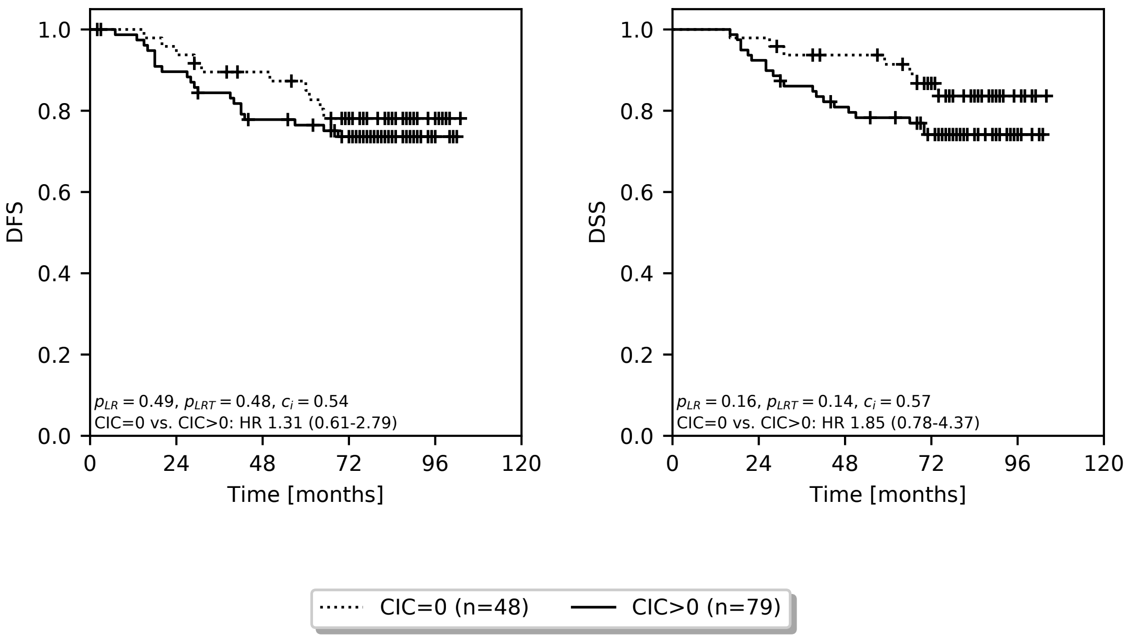

**Fig. 6G.** Kaplan-Meier estimates of DFS and DSS comparing stage II patients from the NI240 cohort grouped by the absence or presence of CIC events detected in the invasive front tissue. Patients were classified as CIC-negative (CIC=0) or CIC-positive (CIC>0) if the median number of CIC events detected across multiple biological replicas (TMA sections) prepared from the invasive front tissue was equal to or greater than zero, respectively. Kaplan-Meier estimators, logrank p-values and univariate Cox regression models were computed with the python package *lifelines*.

#### Figure 6 panel H

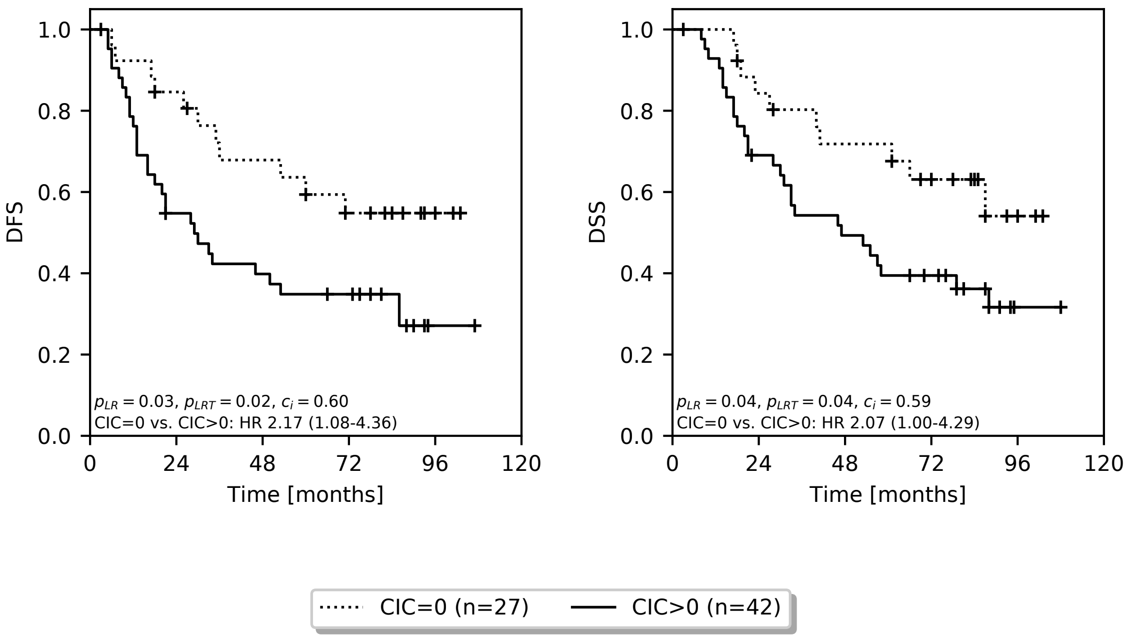

**Fig. 6H.** Kaplan-Meier estimates of DFS and DSS comparing stage III patients from the NI240 cohort grouped by the absence or presence of CIC events detected in the invasive front tissue. Patients were classified as CIC-negative (CIC=0) or CIC-positive (CIC>0) if the median number of CIC events detected across multiple biological replicas (TMA sections) prepared from the invasive front tissue was equal to or greater than zero, respectively. Kaplan-Meier estimators, logrank p-values and univariate Cox regression models were computed with the python package *lifelines*.

### Supplementary Figures

#### Supplementary Figure 1

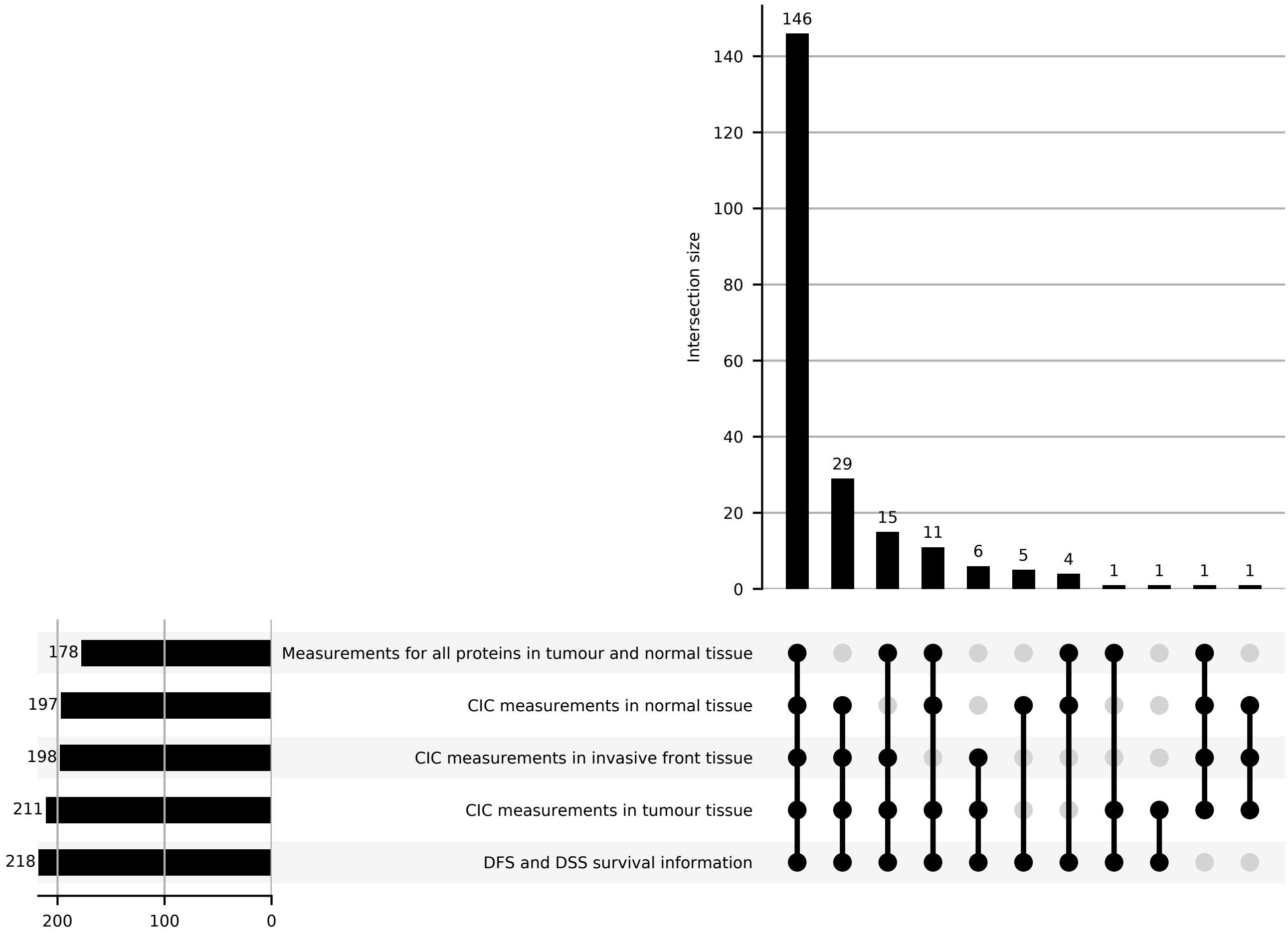

**Sup. Fig. 1.** Data availability per-patient. Figure panel indicating the total number of patients available for analysis of the association between the number of CIC events detected by tissue type (tumour, invasive front and normal), clinical outcome (DFS and DSS) and IHC measurements of TRAIL signalling proteins in tumour and normal tissue. Visualization was created with the python package *upsetplot*.

#### Supplementary Figure 2

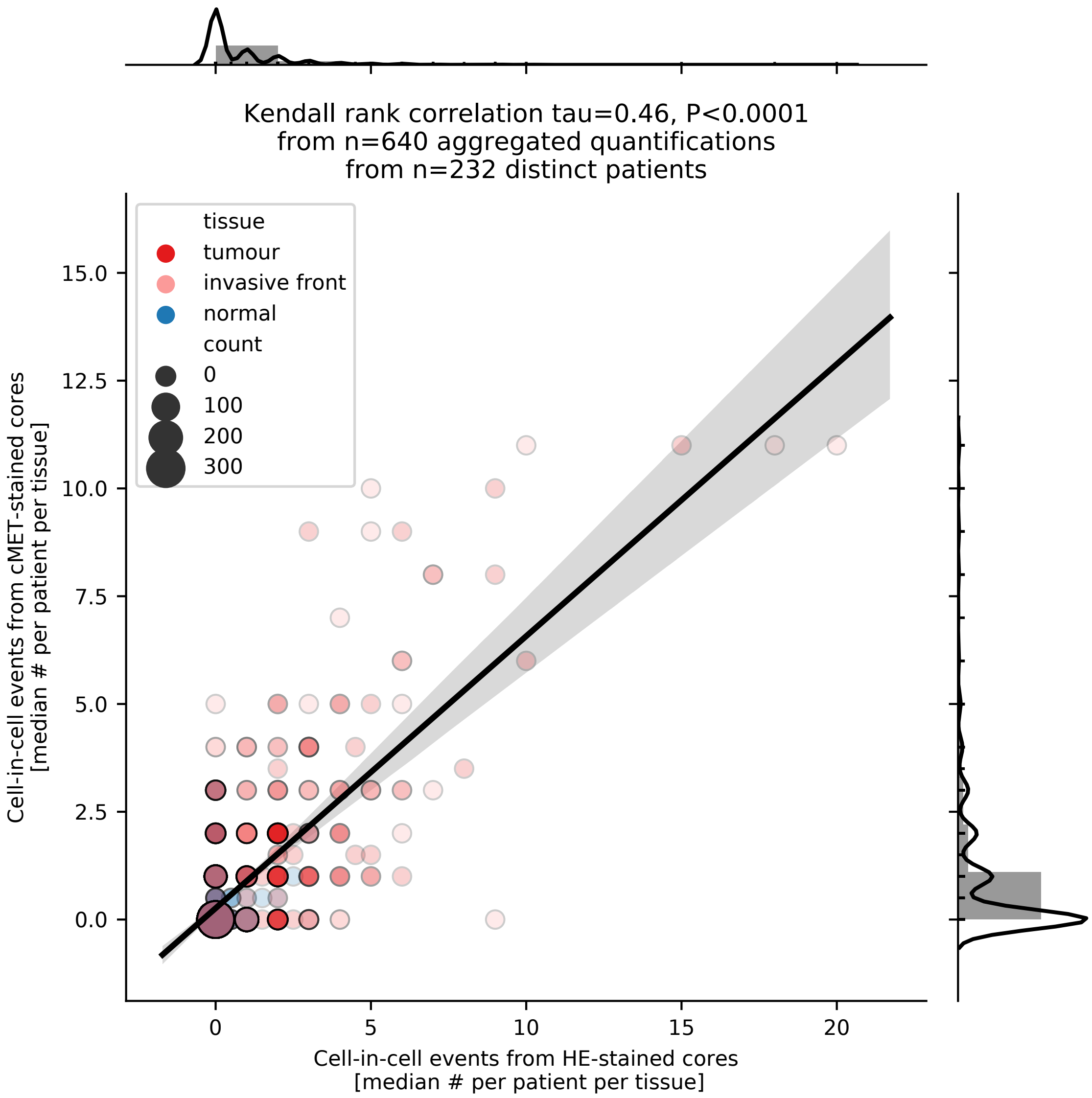

**Sup. Fig. 2.** Comparison between CIC events (median aggregated by patient, tissue and staining marker) observed in TMA sections stained for HE and cMET. Marker color indicates tissue type. Marker size and transparency encode the number of cores. Solid black line and gray shaded area indicate the regression line and CI, respectively. Agreement between CIC estimates from HE- and cMET-stained TMA cores was computed using the Kendall tau correlation (python package *scipy*).

#### Supplementary Figure 3

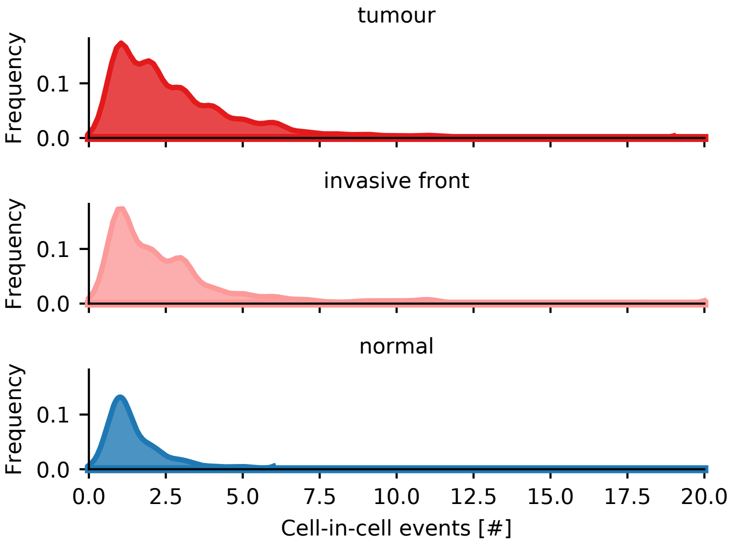

**Sup. Fig. 3.** Distribution of CIC events detected across all TMA sections examined (**Sup. Tables 3-4**) grouped by tissue type.

#### Supplementary Figure 4

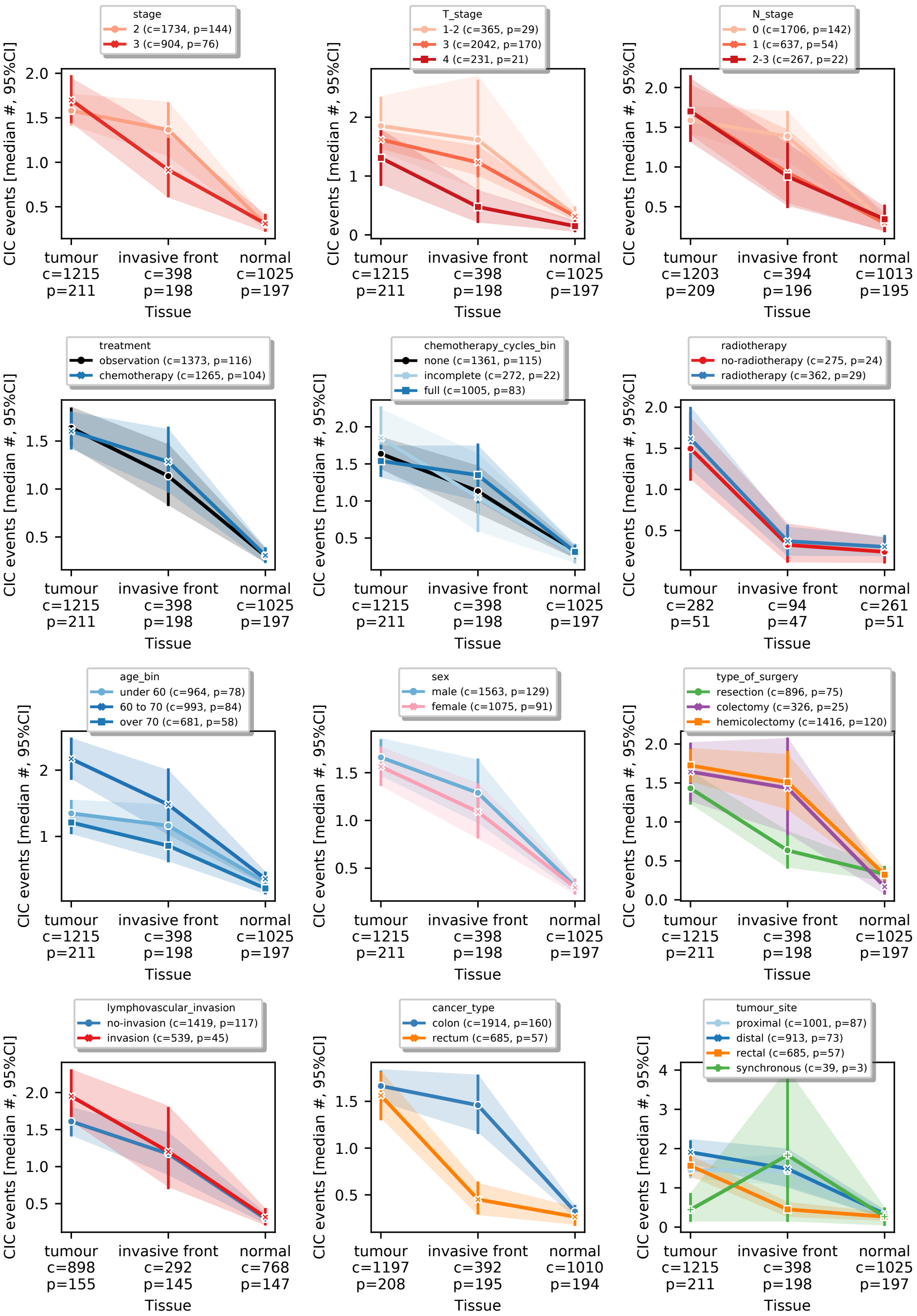

**Sup. Fig. 4.** CIC events (median and 95% CI) detected in TMA sections prepared from tumour centre, invasive front and matched normal tissue broken down by clinico-pathological characteristics. The letters “c” and “p” are abbreviations for “cores” and “patients”, respectively. Corresponding statistical analysis is reported in **Sup. Table 7**.

#### Supplementary Figure 5

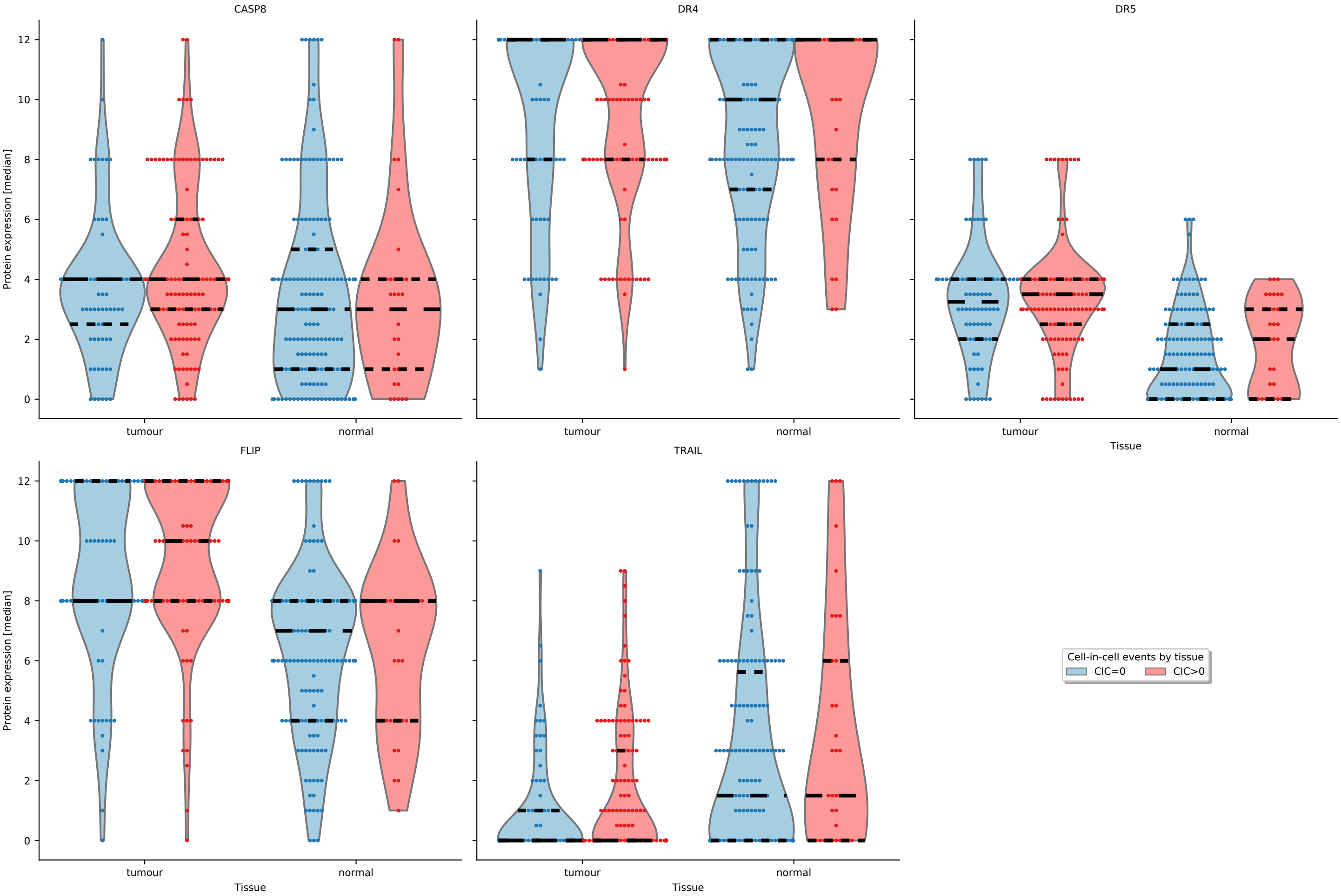

**Sup. Fig. 5.** Association between expression of key proteins involved in TRAIL signalling and CIC events. Protein expression was determined by IHC in TMA sections prepared from tumour and normal tissue and expressed as the product of the staining scores for intensity (0-3 integer scale) and coverage (0-4 integer scale). Protein expression across multiple biological replicas (cores) per patient per tissue type was aggregated by median and plotted grouped by absence (CIC=0) or presence (CIC>0) of CIC events in the corresponding tissue. Protein expression by tissue type and color-coded by CIC events group is shown as violin plot overlaid with individual measurements shown as swarmplot. Distribution quartiles are highlighted by tick dotted black lines. Corresponding statistical analysis (restricted to tumour tissue) is reported in **Sup. Table 8**.

### Supplementary Figure 6

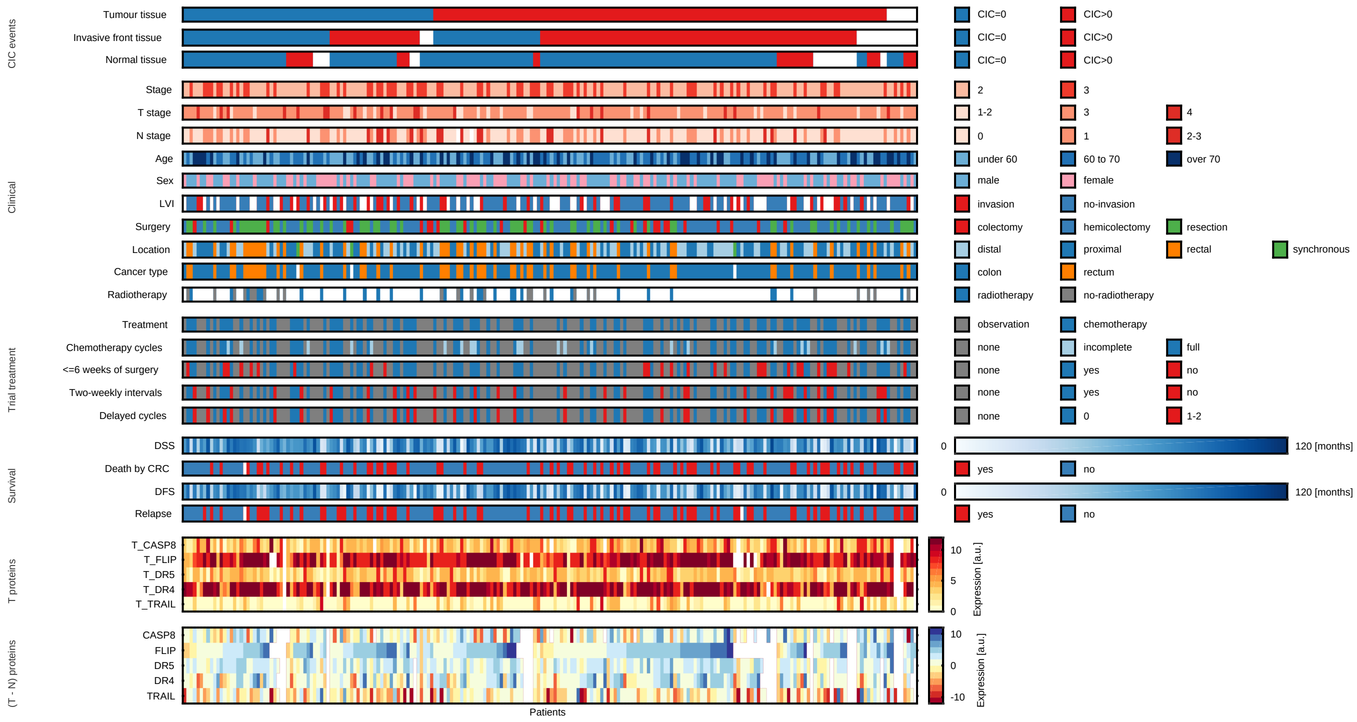

**Sup. Fig. 6.** Overview of clinical, demographic, pathological and molecular information for the CRC patients of the NI240 phase III clinical trial. Each column represents a patient and each row color-codes a feature. Missing data are shown in white. CIC-derived features include estimates of absence (CIC=0) or presence (CIC>0) of CIC events from TMA sections prepared from tumour, invasive front and normal tissue. Visualization was generated with the MATLAB package *HCP*.

#### Supplementary Figure 7

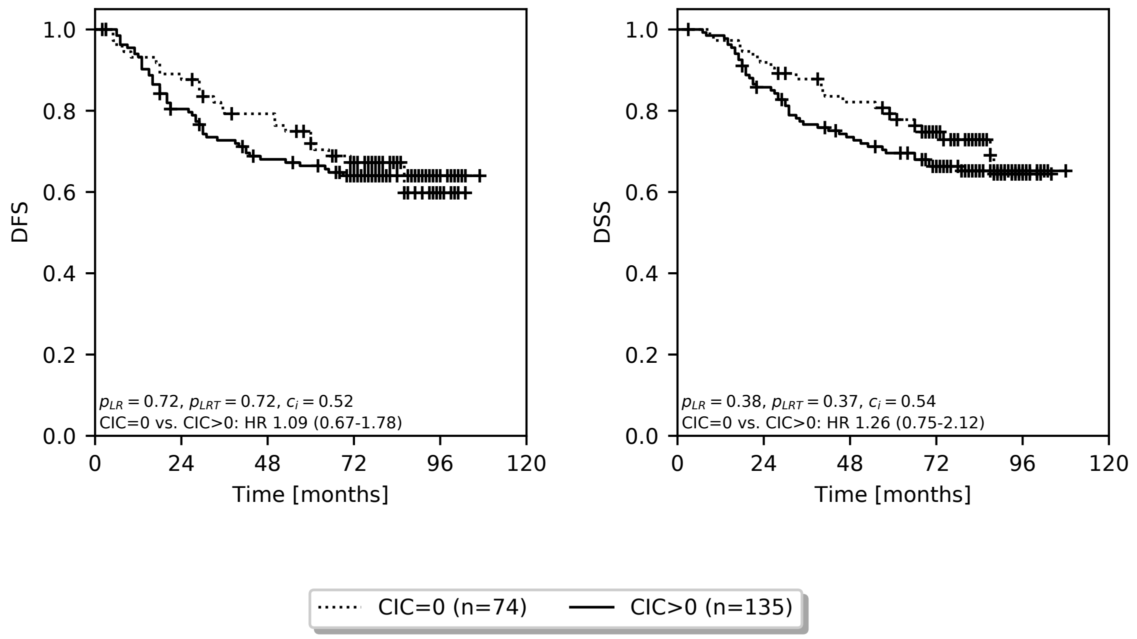

**Sup. Fig. 7.** Kaplan-Meier estimates of DFS and DSS comparing stage II and III patients of the NI240 cohort grouped by the absence or presence of CIC events detected in tumour tissue. Patients were classified as CIC-negative (CIC=0) or CIC-positive (CIC>0) if the median number of CIC events detected across multiple biological replicas (TMA sections) prepared from tumour tissue was equal to or greater than zero, respectively. Kaplan-Meier estimators, logrank p-values and univariate Cox regression models were computed with the python package *lifelines*.

#### Supplementary Figure 8

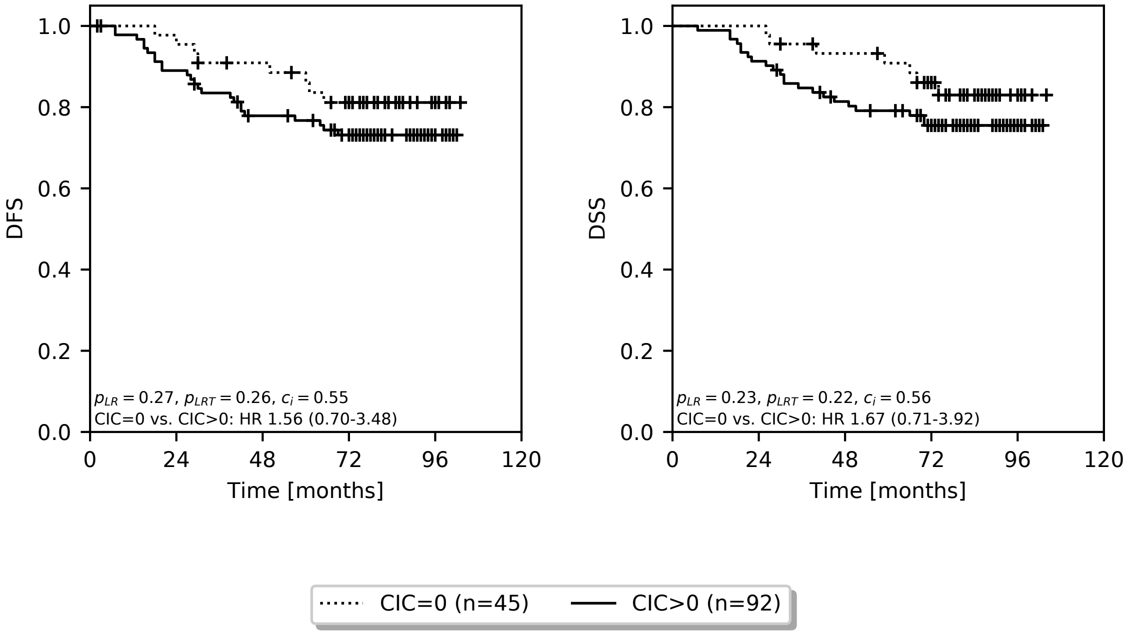

**Sup. Fig. 8.** Kaplan-Meier estimates of DFS and DSS comparing stage II patients of the NI240 cohort grouped by the absence or presence of CIC events detected in tumour tissue. Patients were classified as CIC-negative (CIC=0) or CIC-positive (CIC>0) if the median number of CIC events detected across multiple biological replicas (TMA sections) prepared from tumour tissue was equal to or greater than zero, respectively. Kaplan-Meier estimators, logrank p-values and univariate Cox regression models were computed with the python package *lifelines*.

#### Supplementary Figure 9

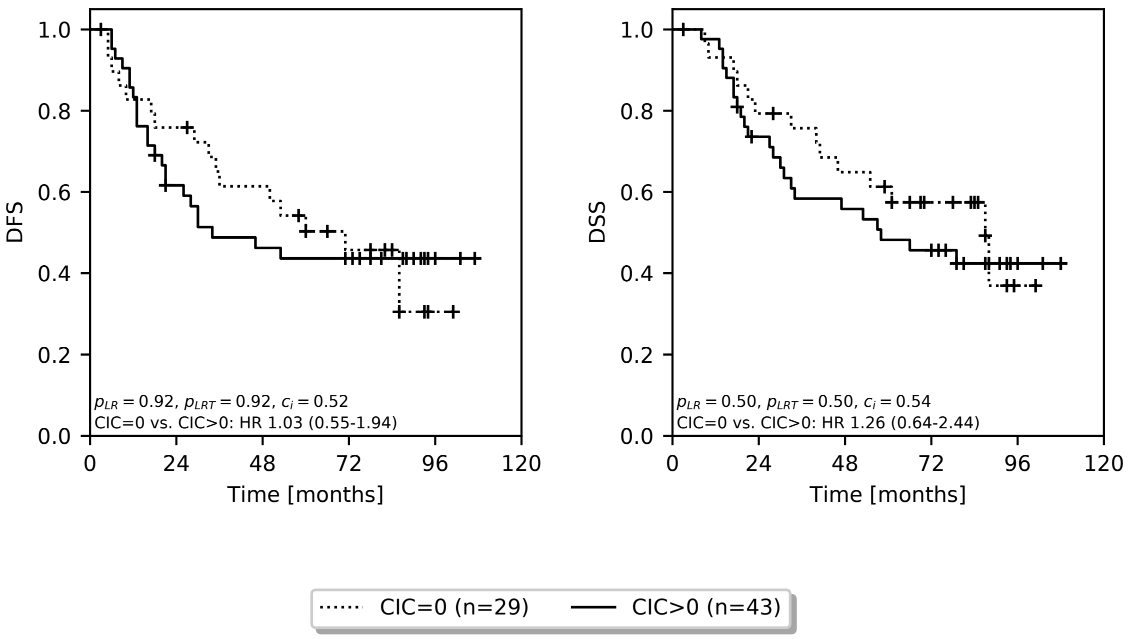

**Sup. Fig. 9.** Kaplan-Meier estimates of DFS and DSS comparing stage III patients of the NI240 cohort grouped by the absence or presence of CIC events detected in tumour tissue. Patients were classified as CIC-negative (CIC=0) or CIC-positive (CIC>0) if the median number of CIC events detected across multiple biological replicas (TMA sections) prepared from tumour tissue was equal to or greater than zero, respectively. Kaplan-Meier estimators, logrank p-values and univariate Cox regression models were computed with the python package *lifelines*.

### Supplementary Tables

#### Supplementary Table 1

**Sup. Table 1:** Clinical, demographic and pathological characteristics of the patients of the NI240 phase III clinical trial. Summary statistics table was created with the python package *tableone*.

| Variable | Level | Missing [n] | Chemotherapy | Observation | P-value | Test |
|---|---|---|---|---|---|---|
| n | | | 117 | 122 | | |
| radiotherapy | no-radiotherapy | 179 | 10 (36) | 17 (53) | 0.275 | Chi-squared |
| | radiotherapy | | 18 (64) | 15 (47) | | |
| cancer type | colon | 4 | 83 (73) | 88 (72) | 0.936 | Chi-squared |
| | rectum | | 30 (27) | 34 (28) | | |
| type of surgery | colectomy | 0 | 9 (8) | 17 (14) | 0.181 | Chi-squared |
| | hemicolectomy | | 69 (59) | 60 (49) | | |
| | resection | | 39 (33) | 45 (37) | | |
| stage | 2 | 0 | 77 (66) | 79 (65) | 0.971 | Chi-squared |
| | 3 | | 40 (34) | 43 (35) | | |
| T stage | 1-2 | 0 | 15 (13) | 18 (15) | 0.706 | Chi-squared |
| | 3 | | 89 (76) | 94 (77) | | |
| | 4 | | 13 (11) | 10 (8) | | |
| N stage | 0 | 2 | 75 (65) | 79 (65) | 0.975 | Chi-squared |
| | 1 | | 28 (24) | 31 (25) | | |
| | 2-3 | | 12 (10) | 12 (10) | | |
| grade | 1 | 10 | 9 (8) | 10 (9) | 0.889 | Chi-squared |
| | 2 | | 88 (78) | 92 (79) | | |
| | 3 | | 16 (14) | 14 (12) | | |
| age | | 0 | 65 [58,70] | 65 [56,72] | 0.958 | Kruskal-Wallis |
| sex | female | 0 | 50 (43) | 46 (38) | 0.509 | Chi-squared |
| | male | | 67 (57) | 76 (62) | | |
| lymphovascular invasion | invasion | 63 | 25 (29) | 23 (26) | 0.794 | Chi-squared |
| | no-invasion | | 62 (71) | 66 (74) | | |

#### Supplementary Table 2

**Sup. Table 2:** Clinical follow-up computed with the python package *lifelines* for the patients of the NI240 phase III clinical trial.

| Endpoint | # patients | Median [months] | 2.5% CI [months] | 97.5% CI [months] |
|---|---|---|---|---|
| DFS | 237 | 82 | 80 | 84 |
| DSS | 238 | 83 | 80 | 85 |

#### Supplementary Table 3

**Sup. Table 3:** Breakdown of the number of TMA cores analyzed for CIC events per patient per tissue from n=232 distinct patients of the NI240 phase III clinical trial.

| Tissue | Total # patients | Total # cores | Median # cores per-patient | Min # cores per-patient |
|---|---:|---:|---:|---:|
| tumour | 223 | 1284 | 6 | 2 |
| invasive front | 209 | 420 | 2 | 2 |
| normal | 209 | 1087 | 6 | 2 |

#### Supplementary Table 4

**Sup. Table 4:** Breakdown of the number of TMA cores analyzed for CIC events per marker, tissue and slide from n=232 distinct patients of the NI240 phase III clinical trial.

| Marker | Tissue | Slide | # patients |
|---|---|---|---|
| HE | tumour | A | 219 |
| | | B | 221 |
| | | C | 208 |
| HE | invasive front | D | 210 |
| HE | normal | E | 187 |
| | | F | 190 |
| | | G | 175 |
| cMET | tumour | A | 210 |
| | | B | 216 |
| | | C | 210 |
| cMET | invasive front | D | 210 |
| cMET | normal | E | 174 |
| | | F | 186 |
| | | G | 175 |

#### Supplementary Table 5

**Sup. Table 5:** Breakdown of the number of TMA cores analyzed for CIC events per patient per tissue for protein expression by IHC for the NI240 phase III clinical trial.

| Tissue | Protein | Total # patients | Total # cores | Median # cores per-patient | Min # cores per-patient |
|---|---|---|---|---|---|
| tumour | CASP8 | 223 | 4150 | 4 | 1 |
| | DR4 | 227 | 4150 | 4 | 1 |
| | DR5 | 229 | 4150 | 4 | 1 |
| | FLIP | 221 | 4150 | 4 | 1 |
| | TRAIL | 230 | 4150 | 4 | 1 |
| normal | CASP8 | 210 | 3451 | 3 | 1 |
| | DR4 | 213 | 3451 | 4 | 1 |
| | DR5 | 217 | 3451 | 4 | 1 |
| | FLIP | 206 | 3451 | 4 | 1 |
| | TRAIL | 218 | 3451 | 4 | 1 |

#### Supplementary Table 6

**Sup. Table 6:** Dependency of the number of observed CIC events detected in TMA sections by tissue type. For statistical analysis, a zero-inflated Poisson regression model, including a random effect for each patient, was fit (R package glmmTMB and car). Number of cores included, effect sizes (estimates), 95% CIs and p-values computed by likelihood ratio tests were reported.

| Term | Ref-level | Level | Estimate | 2.5 % | 97.5 % | P | # cores |
|---|---|---|---|---|---|---|---|
| **tissue** | | | | | | <0.0001 | 606 |
| | **tumour** | **invasive front** | -0.06 | -0.23 | 0.12 | | 606 |
| | | **normal** | -2.36 | -2.78 | -1.94 | | 606 |

#### Supplementary Table 7

**Sup. Table 7:** Association between number of observed CIC events detected in TMA sections and clinical, demographic or pathological covariates by tissue type. Variables were selected *a priori*. For statistical analysis, a zero-inflated Poisson regression model, including a random effect for each patient and covariate-fixed effects, was fitted (R package *glmmTMB* and car). Number of cores included, effect sizes (estimates) for the variable of interest, 95% CIs and p-values computed by likelihood ratio tests were reported.

| Term | Ref-level | Level | Estimate | 2.5 % | 97.5 % | P | # cores |
|---|---|---|---|---|---|---|---|
| **stage** | | | | | | 0.39 | 606 |
| | **2** | **3** | -0.14 | -0.45 | 0.18 | | 606 |
| **T stage** | | | | | | 0.07 | 606 |
| | **1-2** | **3** | -0.01 | -0.45 | 0.43 | | 606 |
| | | **4** | -0.67 | -1.36 | 0.01 | | 606 |
| **N stage** | | | | | | 0.58 | 600 |
| | **0** | **1** | -0.19 | -0.56 | 0.17 | | 600 |
| | | **2-3** | -0.05 | -0.55 | 0.45 | | 600 |
| **age** | | | | | | 0.13 | 606 |
| | **under 60** | **60 to 70** | 0.22 | -0.11 | 0.56 | | 606 |
| | | **over 70** | -0.15 | -0.54 | 0.24 | | 606 |
| **sex** | | | | | | 0.66 | 606 |
| | **male** | **female** | -0.07 | -0.37 | 0.24 | | 606 |
| **type of surgery** | | | | | | 0.06 | 606 |
| | **resection** | **colectomy** | 0.32 | -0.18 | 0.82 | | 606 |
| | | **hemicolectomy** | 0.40 | 0.07 | 0.73 | | 606 |
| **cancer type** | | | | | | 0.009 | 597 |
| | **colon** | **rectum** | -0.47 | -0.82 | -0.12 | | 597 |
| **tumour site** | | | | | | 0.054 | 606 |
| | **proximal** | **distal** | 0.15 | -0.18 | 0.49 | | 606 |
| | | **rectal** | -0.40 | -0.78 | -0.01 | | 606 |
| | | **synchronous** | -0.17 | -1.45 | 1.10 | | 606 |
| **lymphovascular invasion** | | | | | | 0.56 | 447 |
| | **no-invasion** | **invasion** | 0.11 | -0.27 | 0.49 | | 447 |
| **treatment** | | | | | | 0.49 | 606 |
| | **observation** | **chemotherapy** | 0.11 | -0.19 | 0.40 | | 606 |
| **radiotherapy** | | | | | | 0.53 | 149 |
| | **no-radiotherapy** | **radiotherapy** | 0.21 | -0.45 | 0.87 | | 149 |
| **chemotherapy cycles** | | | | | | 0.71 | 606 |
| | **none** | **full** | 0.09 | -0.23 | 0.41 | | 606 |
| | | **incomplete** | 0.19 | -0.32 | 0.69 | | 606 |

#### Supplementary Table 8

**Sup. Table 8:** Association between expression of proteins involved in TRAIL signalling and CIC events observed in tumour tissue of CRC patients of the NI240 phase III clinical trial. Protein expression determined by IHC across biological replicas of tumour sections (**Sup. Table 5**) and per patient were aggregated by the median. CIC events observed across multiple biological TMA cores prepared from tumour tissue were aggregated by median and binarised into absence (CIC=0) or presence (CIC>0). Statistical analysis was performed with the python package statsmodels by univariate unadjusted linear models and effect sizes, 95% CIs, p-values and number of patients included in each analysis were reported.

| Protein | Unadjusted statistics |
|---|---|
| CASP8 | CIC>0 (ref. CIC=0): 0.66, 95% CI -0.02-1.35, P=0.06, n=216 |
| DR4 | CIC>0 (ref. CIC=0): 0.37, 95% CI -0.44-1.18, P=0.37, n=220 |
| DR5 | CIC>0 (ref. CIC=0): 0.12, 95% CI -0.38-0.62, P=0.65, n=222 |
| FLIP | CIC>0 (ref. CIC=0): 0.78, 95% CI 0.00-1.55, P=0.049, n=213 |
| TRAIL | CIC>0 (ref. CIC=0): 0.59, 95% CI 0.02-1.15, P=0.04, n=222 |

### Supplementary Table 9

**Sup. Table 9:** Univariate and multivariate Cox regression models for stage II and III patients of the NI240 phase III clinical trial. Univariate models were fitted for baseline clinical, demographic and pathological characteristics and features derived from CIC events observed in TMA sections prepared from tumour, invasive front and normal tissue. For CIC events-based features, patients were grouped based on the absence (CIC=0) or presence (CIC>0) of CIC events computed as median across multiple cores for the corresponding tissue. The multivariate model included CIC events feature derived from the invasive front and was adjusted by baseline patient characteristics selected *a priori*. HRs, 95% CIs, p-values computed by loglikelihood ratio tests, *c*-indices and number of included patients were reported. Cox regression models were fitted using the python package *lifelines*.

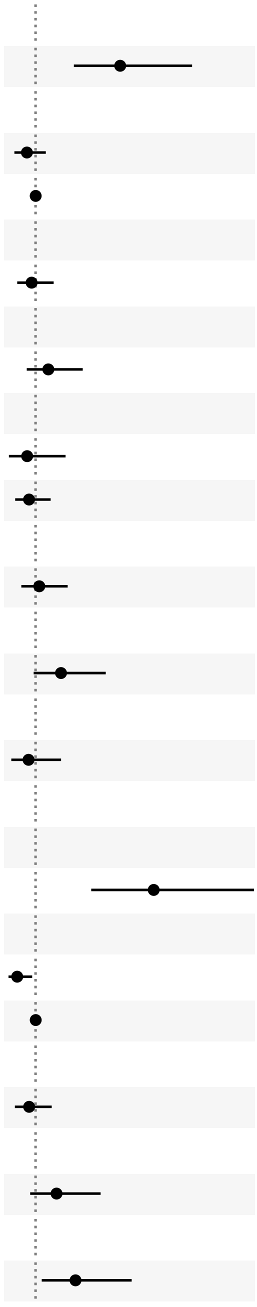

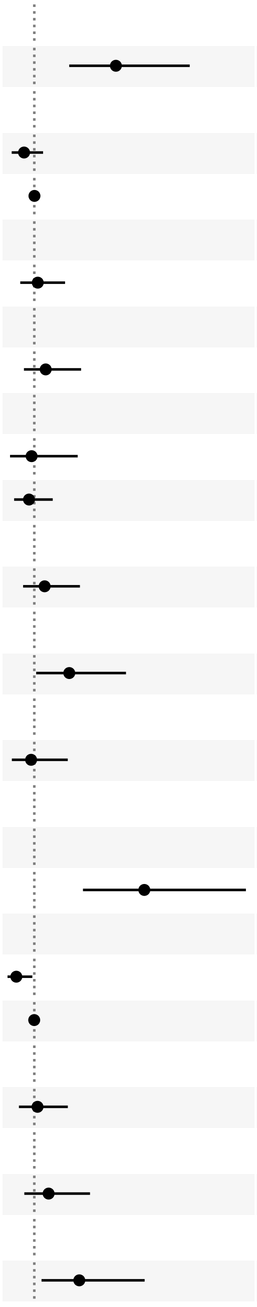

| | | | | DFS | | | | | | | | DSS | | | | | | | |
|---|---|---|---|---|---|---|---|---|---|---|---|---|---|---|---|---|---|---|---|
| Model type | Term | Ref level | Level | HR (95% CI) | HR | 2.5% CI | 97.5% CI | Per-term P-value | Per-model P-value | C-index | N | HR (95% CI) | HR | 2.5% CI | 97.5% CI | Per-term P-value | Per-model P-value | C-index | N |
| univariate | stage | | | | | | | <0.0001 | <0.0001 | 0.64 | 218 | | | | | <0.0001 | <0.0001 | 0.63 | 219 |
| | | 2 | 3 | | 3.13 | 2.00 | 4.91 | | | | | | 3.05 | 1.91 | 4.88 | | | | |
| | treatment | | | | | | | 0.28 | 0.28 | 0.54 | 218 | | | | | 0.21 | 0.21 | 0.54 | 219 |
| | | observation | chemotherapy | | 0.78 | 0.50 | 1.23 | | | | | | 0.74 | 0.46 | 1.19 | | | | |
| | age | | | | 1.00 | 0.98 | 1.03 | 0.82 | 0.82 | 0.52 | 218 | | 1.00 | 0.98 | 1.03 | 0.77 | 0.77 | 0.51 | 219 |
| | sex | | | | | | | 0.65 | 0.65 | 0.51 | 218 | | | | | 0.73 | 0.73 | 0.52 | 219 |
| | | male | female | | 0.90 | 0.57 | 1.42 | | | | | | 1.09 | 0.68 | 1.74 | | | | |
| | cancer type | | | | | | | 0.27 | 0.27 | 0.53 | 215 | | | | | 0.34 | 0.34 | 0.52 | 216 |
| | | colon | rectum | | 1.32 | 0.81 | 2.16 | | | | | | 1.29 | 0.77 | 2.14 | | | | |
| | type of surgery | | | | | | | 0.72 | 0.72 | 0.52 | 218 | | | | | 0.86 | 0.86 | 0.51 | 219 |
| | | resection | colectomy | | 0.79 | 0.36 | 1.72 | | | | | | 0.93 | 0.42 | 2.06 | | | | |
| | | | hemicolectomy | | 0.84 | 0.52 | 1.35 | | | | | | 0.87 | 0.52 | 1.43 | | | | |
| | CIC in tumour tissue | | | | | | | 0.72 | 0.72 | 0.52 | 209 | | | | | 0.38 | 0.38 | 0.54 | 210 |
| | | T-CIC=0 | T-CIC>0 | | 1.09 | 0.67 | 1.78 | | | | | | 1.26 | 0.75 | 2.12 | | | | |
| | CIC in invasive front tissue | | | | | | | 0.051 | 0.051 | 0.56 | 196 | | | | | 0.02 | 0.02 | 0.58 | 197 |
| | | S-CIC=0 | S-CIC>0 | | 1.64 | 0.98 | 2.74 | | | | | | 1.88 | 1.08 | 3.27 | | | | |
| | CIC in normal tissue | | | | | | | 0.56 | 0.56 | 0.51 | 195 | | | | | 0.81 | 0.81 | 0.51 | 196 |
| | | N-CIC=0 | N-CIC>0 | | 0.82 | 0.42 | 1.61 | | | | | | 0.92 | 0.47 | 1.81 | | | | |
| multivariate | | | | | | | | | <0.0001 | 0.71 | 193 | | | | | | <0.0001 | 0.71 | 194 |
| | stage | | | | | | | <0.0001 | | | | | | | | <0.0001 | | | |
| | | 2 | 3 | | 3.98 | 2.43 | 6.50 | | | | | | 3.77 | 2.26 | 6.29 | | | | |
| | treatment | | | | | | | 0.01 | | | | | | | | 0.02 | | | |
| | | observation | chemotherapy | | 0.54 | 0.33 | 0.88 | | | | | | 0.55 | 0.33 | 0.92 | | | | |
| | age | | | | 1.00 | 0.98 | 1.03 | 0.79 | | | | | 1.00 | 0.97 | 1.02 | 0.87 | | | |
| | sex | | | | | | | 0.48 | | | | | | | | 0.77 | | | |
| | | male | female | | 0.84 | 0.51 | 1.38 | | | | | | 1.08 | 0.65 | 1.81 | | | | |
| | cancer type | | | | | | | 0.13 | | | | | | | | 0.29 | | | |
| | | colon | rectum | | 1.53 | 0.90 | 2.61 | | | | | | 1.36 | 0.78 | 2.37 | | | | |
| | CIC in invasive front tissue | | | | | | | 0.007 | | | | | | | | 0.006 | | | |
| | | S-CIC=0 | S-CIC>0 | | 2.01 | 1.19 | 3.39 | | | | | | 2.13 | 1.21 | 3.74 | | | | |

### Supplementary Table 10

**Sup. Table 10:** Univariate and multivariate Cox regression models for stage III patients of the NI240 phase III clinical trial. Univariate models were fitted for baseline clinical, demographic and pathological characteristics and features derived from CIC events observed in TMA sections prepared from tumour, invasive front and normal tissue. For CIC events-based features, patients were grouped based on the absence (CIC=0) or presence (CIC>0) of CIC events computed as median across multiple cores for the corresponding tissue. The multivariate model included CIC events feature derived from the invasive front and was adjusted by baseline patient characteristics selected *a priori*. HRs, 95% CIs, p-values computed by loglikelihood ratio tests, *c*-indices and number of included patients were reported. Cox regression models were fitted using the python package *lifelines*.

| | | | | DFS | | | | | | | | DSS | | | | | | | |
|---|---|---|---|---|---|---|---|---|---|---|---|---|---|---|---|---|---|---|---|
| Model type | Term | Ref level | Level | HR (95% CI) | HR | 2.5% CI | 97.5% CI | Per-term P-value | Per-model P-value | C-index | N | HR (95% CI) | HR | 2.5% CI | 97.5% CI | Per-term P-value | Per-model P-value | C-index | N |
| univariate | treatment | | | | | | | 0.03 | 0.03 | 0.60 | 75 | | | | | 0.11 | 0.11 | 0.57 | 75 |
| | | observation | chemotherapy | | 0.52 | 0.28 | 0.96 | | | | | | 0.60 | 0.31 | 1.13 | | | | |
| | age | | | | 0.99 | 0.96 | 1.03 | 0.77 | 0.77 | 0.49 | 75 | | 0.99 | 0.95 | 1.02 | 0.46 | 0.46 | 0.53 | 75 |
| | sex | | | | | | | 0.80 | 0.80 | 0.51 | 75 | | | | | 0.36 | 0.36 | 0.54 | 75 |
| | | male | female | | 1.08 | 0.58 | 2.01 | | | | | | 1.35 | 0.71 | 2.54 | | | | |
| | cancer type | | | | | | | 0.77 | 0.77 | 0.51 | 75 | | | | | 0.98 | 0.98 | 0.48 | 75 |
| | | colon | rectum | | 1.10 | 0.57 | 2.13 | | | | | | 1.01 | 0.51 | 1.99 | | | | |
| | type of surgery | | | | | | | 0.86 | 0.86 | 0.51 | 75 | | | | | 1.00 | 1.00 | 0.53 | 75 |
| | | resection | colectomy | | 0.79 | 0.23 | 2.71 | | | | | | 1.03 | 0.30 | 3.56 | | | | |
| | | | hemicolectomy | | 0.85 | 0.45 | 1.60 | | | | | | 1.02 | 0.52 | 1.97 | | | | |
| | CIC in tumour tissue | | | | | | | 0.92 | 0.92 | 0.52 | 72 | | | | | 0.50 | 0.50 | 0.54 | 72 |
| | | T-CIC=0 | T-CIC>0 | | 1.03 | 0.55 | 1.94 | | | | | | 1.26 | 0.64 | 2.44 | | | | |
| | CIC in invasive front tissue | | | | | | | 0.02 | 0.02 | 0.60 | 69 | | | | | 0.04 | 0.04 | 0.59 | 69 |
| | | S-CIC=0 | S-CIC>0 | | 2.17 | 1.08 | 4.36 | | | | | | 2.07 | 1.00 | 4.29 | | | | |
| | CIC in normal tissue | | | | | | | 0.50 | 0.50 | 0.52 | 65 | | | | | 0.75 | 0.75 | 0.51 | 65 |
| | | N-CIC=0 | N-CIC>0 | | 0.68 | 0.21 | 2.22 | | | | | | 0.83 | 0.25 | 2.72 | | | | |
| multivariate | | | | | | | | | 0.01 | 0.69 | 69 | | | | | | 0.06 | 0.67 | 69 |
| | treatment | | | | | | | 0.003 | | | | | | | | 0.02 | | | |
| | | observation | chemotherapy | | 0.36 | 0.18 | 0.71 | | | | | | 0.43 | 0.21 | 0.87 | | | | |
| | age | | | | 1.01 | 0.97 | 1.04 | 0.76 | | | | | 0.99 | 0.95 | 1.02 | 0.49 | | | |
| | sex | | | | | | | 0.72 | | | | | | | | 0.28 | | | |
| | | male | female | | 1.13 | 0.59 | 2.18 | | | | | | 1.47 | 0.73 | 2.95 | | | | |
| | cancer type | | | | | | | 0.32 | | | | | | | | 0.83 | | | |
| | | colon | rectum | | 1.45 | 0.70 | 2.97 | | | | | | 1.09 | 0.52 | 2.25 | | | | |
| | CIC in invasive front tissue | | | | | | | 0.004 | | | | | | | | 0.02 | | | |
| | | S-CIC=0 | S-CIC>0 | | 2.67 | 1.31 | 5.43 | | | | | | 2.33 | 1.12 | 4.87 | | | | |

MATLAB. 2014. *Version 8.4.0 (R2014b)*. Natick, Massachusetts: The MathWorks Inc.

McKinney, Wes, and others. 2010. "Data Structures for Statistical Computing in Python." In *Proceedings of the 9th Python in Science Conference*, 445:51–56. Austin, TX.

Oliphant, Travis E. 2006. *A Guide to Numpy*. Vol. 1. Trelgol Publishing USA.

Pollard, Tom J, Alistair E W Johnson, Jesse D Raffa, and Roger G Mark. 2018. "tableone: An open source Python package for producing summary statistics for research papers." *JAMIA Open* 1 (1): 26–31. https://doi.org/10.1093/jamiaopen/ooy012.

R Core Team. 2020. *R: A Language and Environment for Statistical Computing*. Vienna, Austria: R Foundation for Statistical Computing. https://www.R-project.org/.

Salvucci, Manuela, and Jochen H. m. Prehn. 2019. "HCP: A Matlab Package to Create Beautiful Heatmaps with Richly Annotated Covariates." *Journal of Open Source Software* 4 (38): 1291. https://doi.org/10.21105/joss.01291.

Seabold, Skipper, and Josef Perktold. 2010. "Statsmodels: Econometric and Statistical Modeling with Python." In *9th Python in Science Conference*.

Van Rossum, Guido, and Fred L. Drake. 2009. *Python 3 Reference Manual*. Scotts Valley, CA: CreateSpace.

Virtanen, Pauli, Ralf Gommers, Travis E. Oliphant, Matt Haberland, Tyler Reddy, David Cournapeau, Evgeni Burovski, et al. 2020. "SciPy 1.0: Fundamental Algorithms for Scientific Computing in Python." *Nature Methods*. https://doi.org/https://doi.org/10.1038/s41592-019-0686-2.

Waskom, Michael, Olga Botvinnik, Drew O'Kane, Paul Hobson, Saulius Lukauskas, David C Gemperline, Tom Augspurger, et al. 2018. *Mwaskom/Seaborn: V0.9.0 (July 2018)* (version v0.9.0). Zenodo. https://doi.org/10.5281/zenodo.1313201.
